# Bilayer charge asymmetry and oil residues destabilize membranes upon poration

**DOI:** 10.1101/2023.02.03.526964

**Authors:** Fernanda S. C. Leomil, Mareike Stephan, Shreya Pramanik, Karin A. Riske, Rumiana Dimova

## Abstract

Transmembrane asymmetry is ubiquitous in cells, particularly with respect to lipids, where charged lipids are mainly restricted to one monolayer. We investigate the influence of anionic lipid asymmetry on the stability of giant unilamellar vesicles (GUVs), minimal plasma membrane models. To quantify asymmetry, we apply the fluorescence quenching assay, which is often difficult to reproduce and caution in handling the quencher is generally underestimated. Thus, we first optimize this assay and then apply it to GUVs prepared with the inverted emulsion transfer protocol using increasing fractions of anionic lipids restricted to one leaflet. This protocol is found to produce highly asymmetric bilayers, but with ∼20% interleaflet mixing. To probe the stability of asymmetric vs symmetric membranes, we expose the GUVs to porating electric pulses and monitor the fraction of destabilized vesicles. The pulses open macropores, and the GUVs either completely recover or exhibit leakage or bursting/collapse. Residual oil destabilizes porated membranes and destabilization is even more pronounced in asymmetrically charged membranes. This is corroborated by the measured pore edge tension decrease with increasing charge asymmetry. Using GUVs with membranes with imposed transmembrane pH asymmetry, we confirm that poration-triggered destabilization does not depend on the approach used to generate membrane asymmetry.

## Introduction

A typical eukaryotic cell membrane is highly asymmetric in the distribution of its main constituents, which is essential to ensure the distinct functions of cells. The asymmetry is comprehensive in respect to membrane proteins and carbohydrates: integral proteins exhibit always the same orientation, peripheral proteins are only found associated with one of the leaflets and carbohydrates attached to proteins and lipids only face the external medium, where they are crucial to cell signaling. Importantly, the lipid bilayer composition is also highly asymmetric ^1^. Specifically in mammalian membranes, phosphatidylcholine and sphingomyelin are found in abundance in the outer monolayer, while phosphatidylethanolamine and anionic lipids, such as phosphatidylserine and phosphatidylinositol, are most commonly found in the inner leaflet, giving rise to a charge asymmetry across the membrane ^2, 3^. Lipid asymmetry affects many membrane properties, such as curvature, shape, permeability and stability of cell membranes. The loss of this asymmetry implies crucial physiological consequences ^4–7^, as for instance the apoptotic cascade followed by the externalization of phosphatidylserine ^8, 9^. Therefore, healthy cells dedicate substantial effort and energy to sustain membrane asymmetry. This is achieved mainly by the work of flippases and floppases, which are enzymes that transport lipids from one leaflet to the other in order to keep the desired lipid asymmetry ^10,11^, but also via protein-free processes ^12^ (more relevant for model membranes). Reversible lipid asymmetry is now also being recognized as a factor influencing intracellular signaling and intercellular communication ^13^. These efforts highlight the importance of asymmetry and merit in being thoroughly investigated.

Membrane stability is of essential importance to cell viability, as the first collective property of the plasma membrane is to act as a boundary to the cell, regulating the traffic of substances. The integrity of the cell membrane relies mainly on the material properties of the constituting lipid bilayer, which due to the hydrophobic effect forms a cohesive and robust film that is nonetheless soft and able to bend. These properties are sensitive to the lipid composition and are affected by the asymmetric distribution of lipids ^1^, an effect that can be further enhanced when charge asymmetry is present. Even though membranes are stable, they can rupture through the opening of a pore, for example in response to mechanical stress. Poration can lead to cell death in the case that a quick resealing of the pore fails ^14, 15^. Pores in cell membranes can also be created on purpose, with the application of a high-intensity electric pulse ^16^, as in clinical procedures, favoring the entrance of different molecules into cells for which the membrane is generally impermeable ^17^. Due to its efficiency, this method (named electroporation or electropermeabilization) has become a common approach in the treatment of various types of cancer ^18–22^. Additionally, it is being used for gene therapy ^23, 24^ and to encapsulate or promote cargo release in drug delivery systems ^25^.

Model membranes have emerged as a useful tool to allow a better understanding of physiological phenomena involving cell membranes. Being composed of a minimal set of components, they represent a simplified version of complex biomembranes and are less susceptible to possible interferences from different processes. They also offer the benefit of allowing independent changes of a single parameter at a time. In particular, giant unilamellar vesicles (GUVs) ^26^ stand out as an ideal system since they replicate the plasma membrane in terms of size (10 - 100 µm) and curvature and are large enough to be observed and manipulated under an optical microscope. GUVs internal and external aqueous solutions are often chosen to be sucrose and glucose, respectively. In these settings, when observed under phase contrast microscopy, the refractive indices of the two sugar solutions create a contrast across the vesicle membrane making the GUVs interface appear as dark contour with a bright halo around. Furthermore, any discontinuity in the membrane (such as the one caused by the opening of a micron-sized pore) can be easily visualized ^27^. The response of GUVs to electric pulses has been studied in detail, revealing interesting relaxation properties of lipid bilayers ^27, 28^, including pore opening and closing dynamics ^29–31^. It was shown that while electric pulses applied to zwitterionic GUVs composed of POPC (palmitoyl oleoyl phosphatidylcholine) caused the opening of transient macropores that lasted about 50 ms, GUVs containing the anionic lipid POPG (palmitoyl oleoyl phosphatidylglycerol) could be completely disrupted and collapse after the pulse ^32^. The presence of POPG and other anionic lipids and molecules was shown to render the membrane more susceptible to the electric pulses, giving rise to leaky membranes or to complete vesicle collapse (bursting) ^33–35^. A fundamental membrane material property that characterizes the stability of pores formed in a membrane is the pore edge tension (γ), which reflects the energy cost of maintaining an open pore in the membrane ^36^, and is crucial for plasma membrane repair processes. If the cost to rearrange the lipids in the pore rim is too low, as in the case of unstable membranes, the vesicle will burst due to continuous opening of the pore, associated with low edge tension values. The pore edge tension can be measured from the dynamics of macropore closure ^29, 34, 36^. Earlier data have shown a two-fold reduction for membranes containing 50 mol% charged lipids ^32–34^ compared to neutral membranes. Interestingly, the increased destabilization was not observed to depend on the means of poration approach (electric pulse or use of detergent) and on the specific anionic lipid, but rather on the surface charge density in the membrane ^33^.

The above-mentioned studies were performed with symmetric lipid bilayers of giant vesicles prepared mainly by the conventional electroformation method ^37^. Lately, several methods have been developed to allow for the preparation of asymmetric membranes to mimic cell membrane asymmetry. Some of them are based on cyclodextrin-mediated lipid exchange ^38–40^, others (and more abundantly applied to the preparation of GUVs) are based on phase-transfer methods (also known as droplet-transfer or emulsion transfer) ^41–44^ assisted by pipettes/microfluidics ^45–47^ or employing double emulsion templates ^48^; lipid exchange mediated by hemifusion to a supported bilayer has also been applied^49, 50^. However, it is of utmost importance to validate the preparation method by probing the actual membrane asymmetry of the generated vesicles. Different methodologies have been reported so far, including nuclear magnetic resonance (NMR) analysis ^51–54^, neutron reflectometry ^55^, small-angle neutron scattering ^56^ and copper-free click chemistry between outer leaflet lipids and fluorophores ^57^. Monitoring the formation of inward or outward nanotubes upon vesicle deflation also offers a way to infer the presence of asymmetry ^58–61^. While NMR and click chemistry-based techniques are not feasible in every laboratory setup and/or cannot be applied to giant vesicles, spontaneous tubulation offers an easy and straightforward visualization. However, this is not a very quantitative approach as tubulation in vesicles is not amenable to precise characterization and can vary from vesicle to vesicle because of the different area-to-volume values.

An alternative technique is the fluorescence quenching assay, first described in 1991 by McIntyre and Sleight as an assay to determine asymmetric fluorophore distributions in small unilamellar vesicles ^62^. Since this first report, the quenching assay has been ubiquitously applied to access membrane asymmetry in small, large and giant unilamellar vesicles (SUVs, LUVs and GUVs, respectively) or even in living cells, see e.g. ^49, 59, 62–64^. The assay is based on the reduction of NBD (nitrobenzoxadiazol)-labeled lipids by dithionite ions, causing an irreversible inactivation (quenching) of the fluorophore, Figure 1. The majority of the studies employing the quenching assay are performed on suspensions of SUVs or LUVs, where the lipid concentration is in the millimolar range, while in GUV suspensions it is orders of magnitude lower and roughly in the micromolar range. This mismatch implies that protocols across systems cannot be applied without adjustment, for example, to match the quencher-to-lipid ratios. Indeed, verbal exchange with researchers in other groups has suggested that this assay when applied to GUVs is not easy to reproduce from lab to lab and between users, presumably due to different initial conditions such as lipid concentrations and buffers. This is why, here, we thoroughly explored and identified conditions that must be ensured for a functional and reproducible quenching assay.

**Figure 1.**
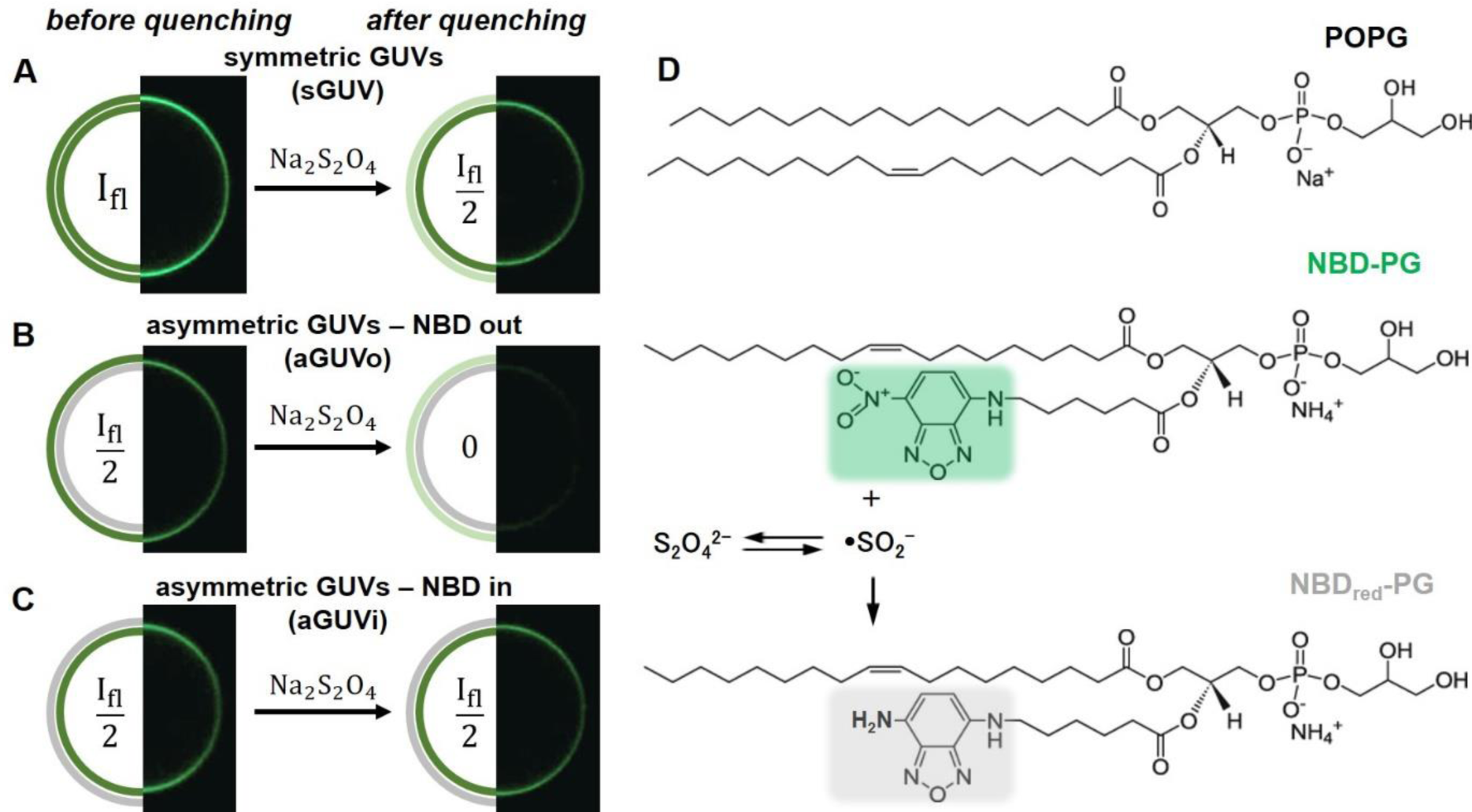
Principle of the quenching assay for evaluating membrane asymmetry in GUVs and chemical structures of molecules. (A-C) The cartoons and example confocal cross sections of GUV halves illustrate how the vesicle fluorescence intensity, I_fl_, should change upon external addition of sodium dithionite when the distribution of the quenched fluorophore in the initial GUV is (A) symmetric (sGUVs) or (B, C) asymmetric with the fluorescent lipid located at the outer or inner leaflet (aGUVo or aGUVi respectively). The vesicles in the shown confocal cross sections had diameters between 20 and 40 μm. (D) Chemical structures of the anionic lipid POPG, its fluorescence analogue NBD-PG and the effect of sodium dithionite on the fluorescent group. In solution, the dithionite ion 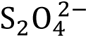 is in equilibrium with the 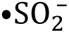 radical, which reduces the nitro group of NBD to its corresponding amine. The reduced NBD-PG (NBD_red_-PG) is non-fluorescent.

Using the optimized assay, we investigate the influence of charge asymmetry in GUVs subjected to electroporation. Asymmetric GUVs composed of POPC with increasing molar fractions of POPG restricted to one of the leaflets are prepared using the inverted emulsion technique. The success of the preparation method and the degree of asymmetry achieved is verified using the quenching of NBD-labeled PG lipid by sodium dithionite. We then interrogate the stability of asymmetric GUVs compared to symmetric ones, by quantifying the fraction of destabilized vesicles upon electroporation and by measuring the pore edge tension, which governs pore closure. We show that membrane destabilization can be much more pronounced if charge asymmetry, as in the case of real cells, is present. Moreover, we raise concerns about possible oil contamination in the membranes of GUVs prepared via the inverted emulsion technique. Finally, an alternative preparation method of asymmetric GUVs, based on pH asymmetry, is put forward to demonstrate that charge asymmetry is the main source of membrane destabilization.

## Materials and Methods

### Materials

The lipids 1-palmitoyl-2-oleoyl-sn-glycero-3-phosphocholine (POPC), 1-palmitoyl-2-oleoyl-sn-glycero-3-phospho-(1’-rac-glycerol) (sodium salt) (POPG) and (1-oleoyl-2-{6-[(7-nitro-2-1,3-benzoxadiazol-4-yl)amino]hexanoyl}-sn-glycero-3-[phospho-rac-(1 glycerol)] (ammonium salt)) (NBD-PG) were purchased from Avanti Polar Lipids (Alabaster, AL). Glucose, sucrose, NaCl, EDTA, TrisHCl (Trizma hydrochloride) and sodium dithionite were purchased from Sigma-Aldrich (St. Louis, MO, USA). Lipids and dye were dissolved in chloroform and the stock solutions were stored at −20 °C until use. Alexa647 hydrazide was purchased from Thermo Fischer (Germany). Light mineral oil was purchased from Carl Roth (Karlsruhe, Germany).

### Vesicle preparation via electroformation

For the optimization of the quenching assay, symmetric GUVs were prepared by the electroformation method ^65^. Briefly, a lipid mixture (6 µL, 2 mM) dissolved in chloroform was spread on the surfaces of two conductive glasses (coated with indium tin oxide), which, after being dried under a stream of nitrogen, were sandwiched, with a Teflon spacer (2 mm thick) forming a chamber (∼2 mL volume). This chamber was filled with sucrose solution (0.2 M) and connected to a function generator. An AC field (1.6 Vpp, 10 Hz) was applied for 30 minutes to accelerate the growth of the GUVs. The vesicles were then harvested and diluted in isotonic glucose solution. The osmolarity was adjusted with an osmometer (Osmomat 3000, Gonotec GmbH, Germany).

### Vesicle preparation via inverted emulsion technique

The protocol for preparation of GUVs via inverted emulsion technique was adapted from previous work on the method ^43, 66^. Briefly, the first step consisted in preparing the lipid-in-oil solutions that were used to create the individual monolayers. The lipid mixture for each monolayer was prepared in a glass vial and the chloroform removed under a stream of nitrogen followed by further evaporation in vacuum for 1h. Mineral oil was added to each vial to give an 800 µM and 400 µM solution for the outer and inner leaflets, respectively. Lipids were dissolved in the oil by sonication for 2h. For the preparation of the outer monolayer, 250 µL of glucose (0.18 or 0.58 M, depending on the experiment) were added to a 1.5 mL protein LoBind tube (Eppendorf, Germany), followed by the addition of 250 µL of lipid-in-oil for the outer leaflet (800 µM), creating a water-oil column. This column was left to stabilize for 2 hours. The next step, after the 2 hours column incubation, consisted in preparing the emulsion of aqueous droplets in oil phase containing the lipids for the inner monolayer. Lipid in oil for the inner leaflet (150 µL, 400 µM) was placed in a separate 1.5 mL tube, followed by the addition of sucrose (4 µL, 0.2 M or 0.6 M, slightly higher osmolarity than the glucose solution). A water-in-oil emulsion was produced by mechanical agitation, dragging 4 times the tube over a tube rack (Polypropylene, 96 positions). The emulsion was then carefully pipetted and deposited on the top of the water-oil column, followed by centrifugation (130g, 10 minutes). After centrifugation, the residual oil on the top of the glucose solution was removed, without extreme perturbation to the interface and the vesicles harvested.

### Imaging of GUVs using optical and confocal microscopy

Different modes of observation were employed. Electroporation experiments for pore edge tension calculation were performed on a Zeiss Axiovert 135 TV (Jena, Germany) phase-contrast inverted microscope equipped with an ultra-fast camera Phantom V2512 (up to 25000 frames per second) or alternatively with an Axio Observer D1 (Jena, Germany) equipped with an sCMOS camera (pco.edge, PCO AG, Kelheim, Germany), for the quantification of GUV response to DC pulse and posterior calculation of the fraction of destabilized vesicles. In both cases, a 20x (NA 0.4) air objective was used. Fluorescence measurements were performed on a Leica confocal SP5 setup (Mannheim, Germany), through a 40x (0.75 NA) air objective. NBD-PG was excited using the 488 nm line of an Argon laser and collected in the 500–600 nm. The Alexa-647 fluorophore was excited using a 638 HeNe laser and the signal was collected between 650-750 nm.

### Python code for measuring membrane fluorescence intensity

The algorithm is provided in the form of Jupyter notebooks, which are files that can be run in a browser. The inputs are “.lif” files, which are the standard file format for Leica confocal microscopes. First, a Gaussian Filter (kernel size of one pixel) was applied to the images to remove noise. To obtain an estimate of the membrane fluorophore concentration, four lines were drawn across the membrane (vertical and horizontal and passing through the GUV center; this approach eliminates contributions form polarization artifacts) to generate four intensity profiles. The integrated area below these intensity profiles is proportional to the fluorophore concentration in the membrane and the mean value of these four measurements was used as mean fluorescence intensity. The code can be found in the GitHub depository: https://github.com/fernandaleomil/fluorescenciaguvs.

### Membrane asymmetry revealed via leaflet specific fluorophore quench

To assess the asymmetric distribution of POPG in the membrane, the dithionite quenching assay was employed targeting NBD-PG. In the first step, the membrane signal of not quenched GUVs was measured by confocal microscopy imaging at the equatorial plane and image analysis using a custom written python code (see previous section). In the next step, the membrane signal of GUVs from a quenched sample was measured and normalized by the mean value obtained on GUVs from the not quenched control sample. At least 20 GUVs were considered for each sample; the scatter in the data results from imaging vesicles of different size (corresponding to different depth in the sample) and inhomogeneity during mixing. The basic procedure of the NBD fluorophore quenching is described in the following (the details for optimizing this protocol are described in the results section on optimization of the quenching assay): freshly prepared sodium dithionite solution (100 mM in 1 M TrisHCl pH 10) was added to a premixed GUV-in-sucrose solution (17.5 µL) and 0.18 M glucose solution (the volume was adjusted to the volume of added dithionite to obtain a final volume of 100 µL) to final sodium dithionite concentrations of 0.5, 1, 1.5, 2, 2.5 or 10 mM. After a certain incubation time (1, 5, 10 or 15 minutes) the sample was diluted 5-fold with 0.18 M glucose (400 µL), in order to reduce the concentration of sodium dithionite. 100 µL of the sample were used for observation. To optimize the protocol, different sodium dithionite concentrations and incubation times were tested.

### Electroporation experiments

GUVs prepared in sucrose were diluted ∼10-fold in glucose solution (at the same solution osmolarity used for GUV preparation) containing the appropriate additive (NaCl and/or EDTA) and placed in an electroporation chamber (Eppendorf, Hamburg, Germany). The chamber consists of two parallel cylindrical platinum electrodes (92 µm in radius) and 500 µm apart (gap distance)^27^. The chamber was connected to a Multiporator (Eppendorf) for DC electric pulse application (3 kV/cm, 150 µs). Experiments to quantify the number of GUVs that underwent bursting or contrast loss (relative to all vesicles in the field, see also Figure S3) after the DC pulse were performed in glucose (0.58 M) and, if not otherwise indicated, without any additive. Image sequences were typically acquired at 1760 px × 2160 px, with acquisition rate of 10 frames per second, for 5 minutes. Pore edge tension experiments were performed on vesicles grown in the presence of NaCl (0.18 M sucrose and 0.5 mM NaCl) and diluted in glucose to induce oblate deformation during the pulse. Image sequences were typically acquired at 512 px × 512 px with acquisition rates between 3000 to 20000 frames per second. Pore dynamics was assessed with the software PoET ^34^, where for the viscosity of the outer solution (η) we used 1.133×10^−3^ Pa·s. The procedures for assessing the number of destabilized vesicles and for measuring the edge tension was repeated several times for each composition, every time on a fresh sample.

### Microfluidic exchange of external GUV solution for probing membrane asymmetry at asymmetric pH

Chip fabrication: The microfluidic device^67^ was prepared using PDMS and glass coverslips. The PDMS and the curing gel were mixed thoroughly in the ratio of 10:1 before degassing in a vacuum chamber. This mixture was poured over the wafer with the microfluidic design cast and baked at 90 °C for 3 h. After cooling, the PDMS was peeled from the wafer and the devices were separated using a sharp blade. To form the inlet and the outlet of the device, holes were punched using a biopsy punch with a plunger system (Kai Medical). The PDMS device and glass coverslips were treated with plasma (Harrick Plasma) for 1 min and pressed together. The whole setup was placed on a hotplate at 80 °C for 30 min.

Experiments on the GUVs: GUVs with the composition POPC:POPG (8:2) doped with 1 mol% NBD-PG were electroformed in sucrose solution. These were diluted in isotonic glucose solution and loaded onto the microfluidic device. The device contained dead-end side channels, in which the GUVs were loaded using a protocol described in detail previously^67^. Briefly, the GUVs were drawn from the reservoir using a syringe pump (Nemesys). The device was oriented vertically to settle the GUVs in the side channels. To exchange the solution outside the GUVs after placing it under the microscope, the reservoir was filled with the new solution and the syringe pump drew out the solution at the rate of 200μL per hour.

## Results and Discussion

Asymmetric GUVs made of POPC and increasing fractions of POPG restricted to one of the leaflets were prepared by the inverted emulsion protocol ^43^. The efficiency of the method in generating asymmetric GUVs was quantified using the assay based on NBD quenching by sodium dithionite. In the following, we first describe the principle of the method and its application to symmetric and asymmetric vesicles (Figure 1). Next, the method is optimized, which is crucial for valid probing of the degree of asymmetry of the obtained GUVs. Then, the stability of asymmetric GUVs was assessed by the application of DC pulses and quantification of destabilization effects and pore edge tension. Finally, the asymmetry in the vesicle bilayer was achieved by exposing the GUV membrane to different pH conditions inside and outside.

### Principle of the quenching assay

The quenching assay is based on the irreversible reduction of the nitro group of the fluorescent probe NBD by the radical 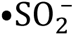 (in equilibrium with the dithionite ion 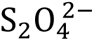) to its corresponding amine, which is non-fluorescent ^62^, see Figure 1D. The fluorescence signal can be quantitatively assessed from confocal microscopy cross sections of GUVs (Figure 1A-C), see also Materials and Methods sections on GUV imaging and fluorescence intensity analysis for details on the image acquisition and processing. For the optimization of the quenching protocol, POPC GUVs with a symmetric distribution of 1 mol% NBD-PG (a PG lipid labeled with NBD in one of the hydrophobic tails, see Figure 1D) were prepared by the conventional electroformation method. Due to the structural and charge similarities, it is expected that NBD-PG distributes across the membrane in a similar manner as POPG and can be treated as its fluorescent representative. Since the NBD group has a relatively high polarity, it is plausible that the lipid tail kinks allowing sodium dithionite to access the NBD fluorophore. Because the membrane is ideally impermeable to sodium dithionite, quenching is expected to only affect fluorophores exposed at the outer membrane leaflet. Therefore, a reduction of 50% in the fluorescence intensity of the membrane is expected due to the symmetric fluorophore distribution, see Figure 1A. For asymmetric vesicles with NBD-PG located only on the outer leaflet (aGUVo), the fluorescence signal should be completely quenched (Figure 1B), while for vesicles with NBD-PG located on the inner leaflet (aGUVi), no change is expected (Figure 1C). The assay requires precaution, since sodium dithionite is a strong reducing agent and unstable in aqueous solutions ^68^. Depending on concentration, pH and oxygen access, dithionite shows different reactions ^69^. Consequently, the optimization of substance handling and reaction conditions are crucial.

### Optimization of the quenching assay

The first concern is to define the best conditions for the preparation of sodium dithionite stock solution. In aqueous environment and in aerobic conditions, sodium dithionite (Na_2_S_2_O_4_) is oxidized to hydrogen sulphide (NaHSO_3_) and hydrogen sulphate (NaHSO_4_) causing a decrease of the solution pH ^70^, which then accelerates further dithionite auto-oxidation. Additional to increased acidity, high dithionite concentration accelerates the decomposition of the dithionite ion^68^. Therefore, the dithionite stock concentration was kept at maximum 0.1 M and not 1 M as described in other publications ^49, 59, 62^. Highly concentrated 1 M sodium dithionite solutions showed yellow color and a strong sulfuric smell, indicating the formation of sulfur dioxide and sulfur. Stock solution concentrations lower than 0.1 M were also avoided to ensure that the volume of the dithionite solution added to the GUV sample is sufficient but small, preventing excessive vesicle dilution. The 0.1 M sodium dithionite solution was always freshly prepared and immediately used.

Previous reports show that alkaline pH-solutions stabilize the dithionite ion ^62, 68, 69^. Therefore, we prepared the 0.1 M sodium dithionite stock solution in 1 M TrisHCl buffer at pH 10. Upon dilution into the vesicle suspension, the solution reached neutral pH. In fact, when the 0.1 M sodium dithionite stock solution was prepared in non-buffered 0.18 M glucose (neutral pH), which is the external solution of the GUVs, the pH of the vesicle solution dropped to 2. Figure S1 shows the importance of preparing the sodium dithionite stock solution at high pH. Whereas the addition of the non-buffered sodium dithionite solution to a GUV sample resulted in permeabilized and defective GUVs and lipid aggregates (Figure S1A), GUVs were preserved when the 0.1 M sodium dithionite stock solution was prepared at pH 10. Efficient quenching with buffered high pH stock solution is exemplified in Figure S1B.

Other important parameters for the quenching assay are the working concentration of sodium dithionite and the incubation time. Note that different concentrations and incubation times have been implemented in the literature (most often using 10 mM final concentration prepared from 1M stock solutions, which, as indicated above, results in vesicle decomposition). Presumably, an additional adjustment of the dithionite concentration is required in the individual working condition, especially if very different total lipid concentrations are explored. It should be stressed, for instance, that lipid, and therefore NBD, concentration in GUV experiments are usually orders of magnitude lower than typical lipid concentrations of LUV/SUV suspensions, for which the quenching assay was originally developed. Here, starting with the stock solution of 0.1 M Na_2_S_2_O_4_ in 1 M TrisHCl pH 10 we tested different final working concentration (from 0.5 to 10 mM Na_2_S_2_O_4_) and incubation times (from 1 to 15 minutes). The desired sodium dithionite concentration was added to the test tube containing the GUVs and after a specific incubation time, the suspension was further diluted 5-fold in order to decrease substantially the quencher concentration, reduce quenching rate, stop unwanted sample degradation and allow for observation and image acquisition (see Figure 2A that demonstrates the effect of quenching the fluorescence signal in both leaflets if the dilution step is not implemented). The results from exploring different incubation times (while implementing the 5-fold dilution step afterwards) are shown in Figure 2B,C. While incubation of 1 minute was not sufficient to inactivate all fluorophores at the outer leaflet, incubation for 5 minutes led to a quenching of roughly 50 % of the total fluorescence (Figure 2C). Longer incubation times of 10 and 15 minutes resulted in fluorescence reductions by more than 50 %, indicating transmembrane dithionite transfer and quenching of part of the inner leaflet fluorophores. Previous studies revealed that the outer leaflet was quenched in the first 50-70 seconds in the case of SUVs, which are much smaller and highly curved ^62^. Then, transfer of dithionite ions or radicals across the membrane was observed to occur at a slower rate leading to quenching of fluorophores residing in the inner leaflet as well (corroborating our results). The authors suggested that the membrane transfer of dithionite ions and radicals depends on the composition and structure of the observed membrane ^62^. Therefore, in the following experiments, GUV samples were diluted to low dithionite concentrations 5 minutes after addition of the quenching agent.

**Figure 2.**
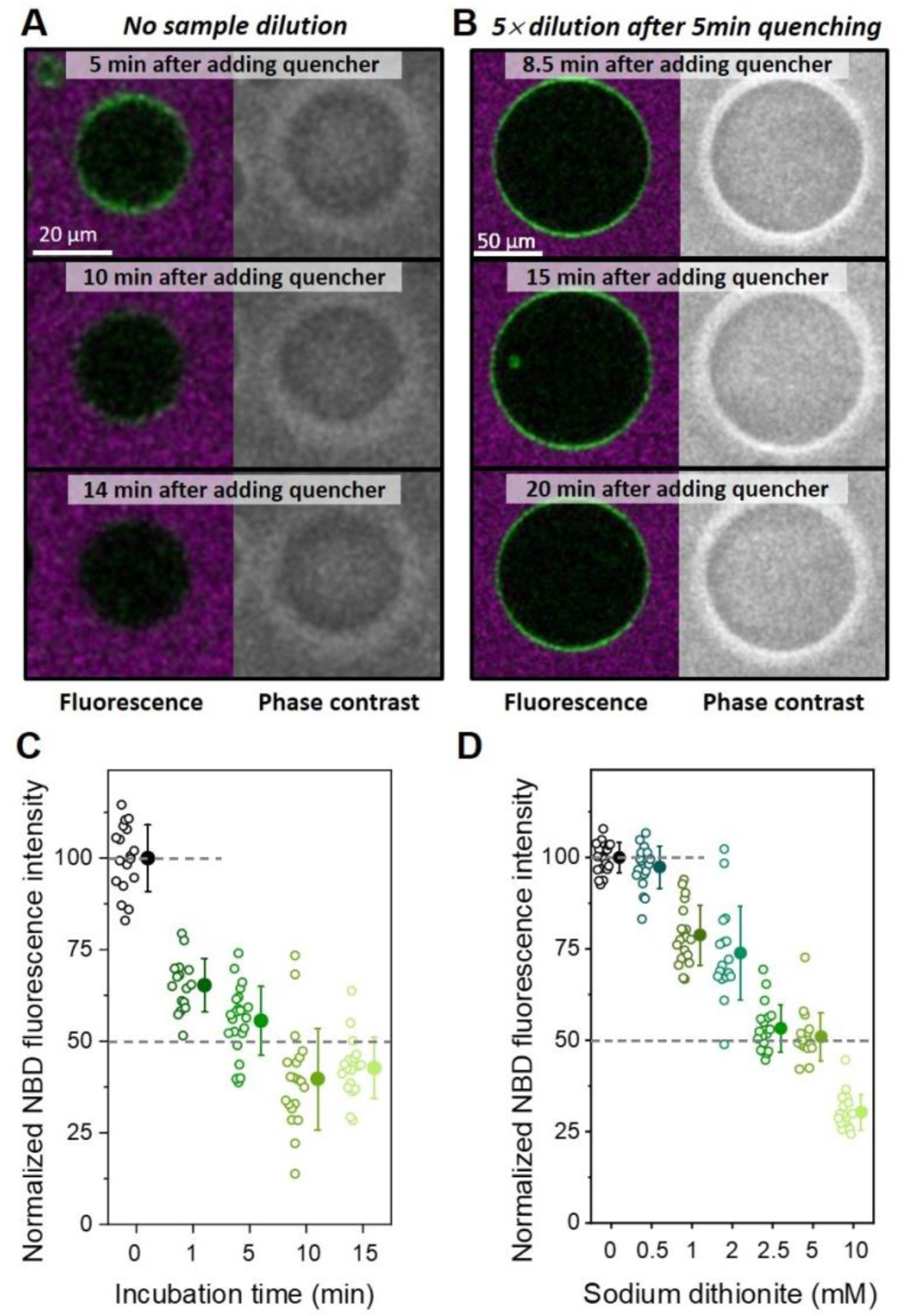
Effect of sample dilution, incubation time and sodium dithionite concentration on the quenching of the NBD fluorescence in GUVs. (A, B) Confocal and phase-contrast image sequences of two symmetric GUVs composed of POPC with 1 mol% NBD-PG (green) prepared with sucrose and then dispersed in glucose medium containing 10 μM of the water-soluble fluorescent dye Alexa 647 (purple). The quenching agent was added to a final concentration of 2.5 mM Na_2_S_2_O_4_ and 5 min after incubation the sample was either directly transferred to the observation chamber (A) or 5-fold diluted in glucose and then transferred for observation (B). Both GUVs remain impermeable to the dye Alexa 647 and sucrose/glucose (optical contrast is maintained), but dithionite ions are able to quench NBD in both leaflets after 14 min in the absence of a dilution step (A). The dilution step restricts quenching to the outer leaflet only (B). (C) Normalized fluorescence intensities before (not quenched, indicated as 0 incubation time) and after quenching with 2.5 mM Na_2_S_2_O_4_ for different incubation times followed by 5-fold dilution. (D) Normalized fluorescence intensities before (0) and after quenching of different sodium dithionite concentration for 5 minutes incubation time followed by 5-fold dilution. Each open symbol represents measurements on one vesicle and mean values with standard deviation are shown as solid symbols on the right. Typically, between 15 and 20 vesicles were measured per sample. The dashed lines are guides to the eye and indicate not-quenched and half of that mean value. Each condition was tested with multiple quenching experiments (n ≥ 3). Note that the x-axis in panels C and D are not linear (see Figure S1 for data presentation with linear x-axis) and that the data are slightly shifted to display individual data points and mean values with SD.

Next, different dithionite concentrations in the final quenching sample were tested. Figure 2D shows that complete quenching of the outer leaflet of GUVs is observed already at final concentration of 2.5 mM. Using higher concentration (10 mM) resulted in fluorescence drops by more than 50 %, indicating that dithionite ions reached some of the inner leaflet fluorophores. We hypothesize that the presence of excess dithionite ions leads to the formation of more decomposition products, which destabilize the membrane and make it permeable for non-decomposed dithionite ions that can then quench the inner leaflet fluorophores. The minimum concentration needed to quench outer leaflet fluorophores depends on the number of fluorophores and on the number of GUVs present in the sample. Therefore, the optimal dithionite concentration for a particular sample should always be tested prior to the actual experiments. For our experimental conditions (roughly 10 μM final total lipid concentration), we chose to work with 2.5 mM dithionite and 5 min incubation time, which was sufficient to quench the NBD groups present only in the external leaflet without causing significant alterations in GUV integrity. It is important to point out how sensitive the quenching assay is to quencher concentration and incubation time (see Figures 2 and S2). Indeed, these conditions should also vary for different lipid concentration. Therefore, it is of utmost importance that these parameters be tested carefully for each working condition before proceeding to obtaining data with the quenching assay.

### Membrane asymmetry of GUVs prepared via the inverted emulsion protocol

In the previous section, we determined important parameters to optimize the quenching assay for probing the membrane asymmetry. We then used the inverted emulsion protocol to obtain POPC GUVs containing 5 mol% POPG in total and 0.5 mol% NBD-PG. The PG lipids were distributed either symmetrically (referred to as sGUVs, with 5 mol% POPG in each membrane leaflet) or asymmetrically (aGUVs, with 10 mol% POPG in one of the leaflets). Two types of asymmetric GUVs were prepared: with POPG and NBD-PG restricted either to the inner (aGUVi) or outer (aGUVo) leaflet. Since we now have the control that sodium dithionite quenches about half of the NBD-PG dyes in sGUVs (Figure 2), we expect either full quenching when the NBD is present only in the outer layer (aGUVo) or no quenching at all if the NBD is restricted to the inner leaflet (aGUVi), as illustrated in Figure 1B,C. Any deviations from these outcomes would imply that lipids from the water-in-oil phase used to form the inner vesicle leaflet have migrated (diffused) and inserted into the oil-water interface with lipids forming the outer GUV leaflet.

The normalized fluorescence intensity before and after quenching of sGUVs and aGUVs grown by the inverted emulsion protocol are shown in Figure 3. As expected, sGUVs have their fluorescence intensity decreased by 50% after addition of sodium dithionite, consistent with data for sGUVs produced via electroformation, thus demonstrating that the employed inverted emulsion protocol efficiently produces GUVs with symmetric distribution of the charged lipids. When aGUVi were exposed to the quenching agent, a signal reduction of ∼25% was observed rather than the expected zero fluorescence intensity reduction. In the case of aGUVo, instead of a complete quenching of the fluorescence, residual fluorescence intensity was detected (∼15%) after treatment with sodium dithionite. Since under the same experimental conditions the symmetric controls showed the expected reduction by 50%, and the integrity of the aGUVs was maintained (observed by preservation of optical contrast under phase contrast), the observed outcome indicates that the inverted emulsion protocol is not efficient in generating entirely asymmetric membranes. We conclude that some mixing (roughly about 20%) of the lipids originating from the different oil layers during the preparation procedure occurred, see estimates in Figure 3. Nonetheless, the inverted emulsion method ensured generation of membranes with a high degree of asymmetry (the leaflet asymmetry in our GUVs is comparable to that reported by Pautot et al.^41^).

**Figure 3.**
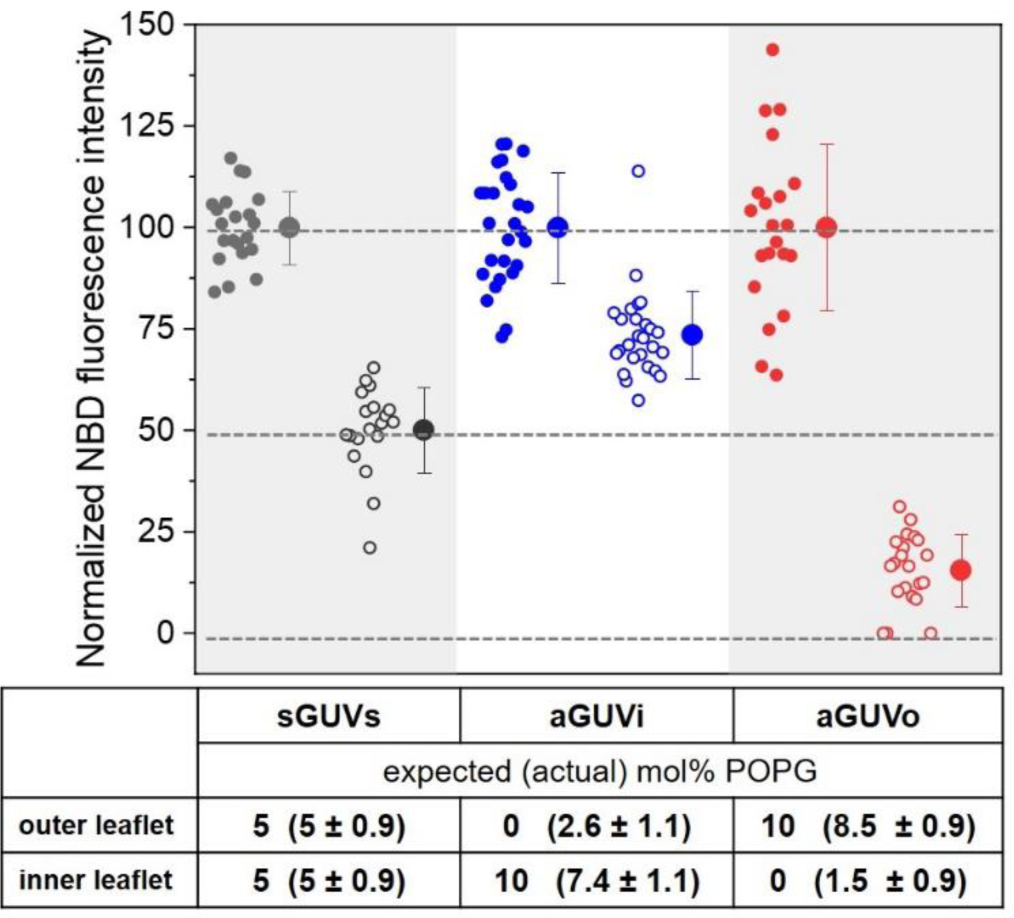
Normalized fluorescence intensities before (solid circles) and after (open circles) quenching of NBD-PG (0.5 mol%) for GUVs containing 5 mol% POPG symmetrically and asymmetrically distributed. Each data point represents measurements on one vesicle and solid circles with error bars on the right indicate mean values with standard deviation. For each type of GUV membrane composition, the fluorescence intensity was normalized by the mean value of the non-quenched measurement. The dashed lines are guides to the eye indicating the mean value of non-quenched GUVs, half of the mean value and zero. All measured GUVs had diameters between 20 and 40 µm. The quenching was done with 2.5 mM Na_2_S_2_O_4_ final concentration, 5 min incubation followed by 5-fold dilution. The table shows the expected molar fraction of POPG in each leaflet as set by the preparation protocol and the one estimated from the fluorescence intensity after quenching the outer leaflet plus the standard deviation. Each condition was tested with multiple quenching experiments and using multiple GUV preparations (n ≥ 3).

### Vesicle stability decreases with increasing membrane charge asymmetry

To assess the effect of membrane asymmetry on GUV stability upon poration, the vesicles were exposed to a single DC pulse (3 kV/cm and 150 μs) and the response was followed with phase contrast optical microscopy. Neutral POPC GUVs typically deformed and the formation of a micrometer-wide pores (macropores) that quickly (∼50 ms) reseal could be observed. Subsequently, the pores reseal and the GUVs restore their integrity with preserved contrast, see Figure 4A. When a similar pulse is applied to symmetric vesicles containing high fractions of anionic lipids, additional effects could occur ^33^. Some GUVs were apparently restored after macropore closure, but remained in a highly permeable state revealed by the loss of sugar asymmetry within 1 minute, indicating that submicroscopic pores persist after the end of the pulse (leaky vesicles, Figure 4B). Still, another fraction of GUVs collapsed, through the indefinite expansion of a macropore, in a phenomenon called bursting (Figure 4C). To quantify the destabilization brought by the presence of charge asymmetry, we applied single DC pulses to a collection of GUVs and evaluated the fraction of vesicles that exhibited any of these two destabilizing effects (leaky state or bursting), see Figure S3 for more information. The fraction of destabilized vesicles (*X_dest_*) was measured for increasing POPG fractions in symmetric and asymmetric GUVs (Figure 4D). The symmetric GUVs were prepared using both electroformation and the inverted emulsion method. Previous studies ^33^ have shown that electroformed and therefore symmetric GUVs are destabilized only at high POPG fractions > 40 mol%. Surprisingly, here we observe that symmetric GUVs prepared by the inverted emulsion method showed considerable membrane destabilization upon electroporation already for pure POPC membranes (neutral vesicles); compare first data points of black and green traces in Figure 4D. This indicates that GUVs prepared by the inverted emulsion technique are less stable compared to electroformed GUVs with the same membrane composition. Presumably, residual oil in the membrane destabilizes the vesicles upon poration. Indeed, Raman scattering microscopy has confirmed the presence of oil in the membrane of GUVs prepared by the droplet transfer method ^71^. In a previous study^72^ we also investigated whether the inverted emulsion approach produces membranes that exhibit different lipid packing and/or differential stress in the membrane^73^ compared to vesicles prepared with the electroformation method. For this, we examined the vesicle morphology upon deflation and measured lipid diffusion. Vesicle deflation in both samples yielded prolate or multisphere GUVs as expected for vesicles, with sucrose/glucose asymmetry across the membrane^74^. Lipid diffusion was also found unaltered. These measurements are compiled in Figure S4. Apparently, the main difference between the vesicles prepared with the two approaches is the presence of oil which locates between the leaflets without affecting lipid packing and symmetry.

**Figure 4.**
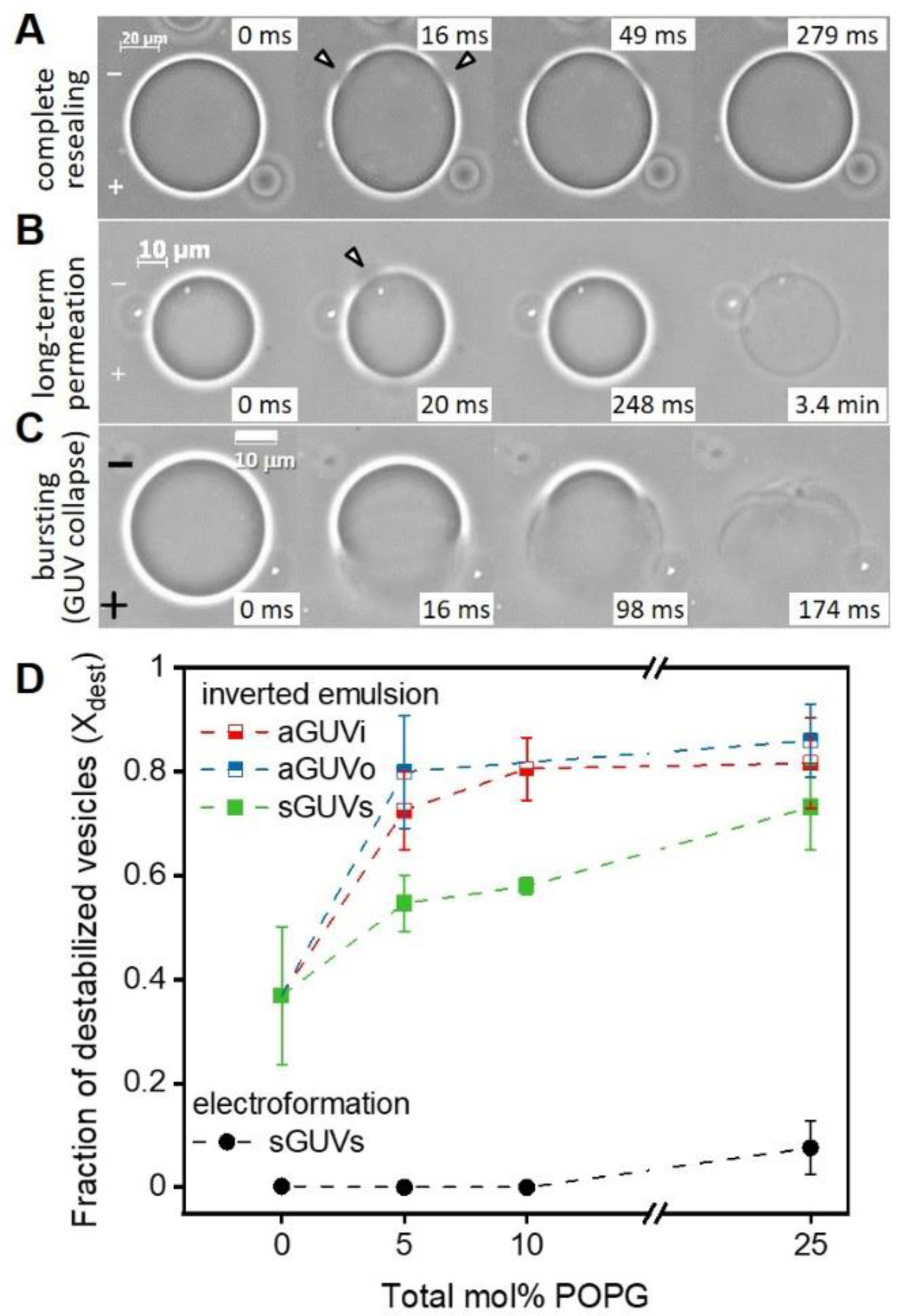
Effect of membrane charge asymmetry on vesicle destabilization upon electroporation. (A-C) Exemplary phase contrast microscopy images showing three possible responses of GUVs to the application of a single DC pulse (3 kV/cm and 150 μs). After the pulse application, micronsized pores (arrowheads) can open and readily reseal restoring GUV integrity (A). Some GUVs that apparently restore their integrity after macropore closure can exhibit high permeability at a later stage revealing the persistence of long-lasting sub-microscopic pores minutes after the end of the pulse (B). In more extreme cases, macropores can open indefinitely leading to vesicle bursting (C). The field polarity is indicated on the first images. (D) Fraction of destabilized vesicles, *X_dest_*, (comprising permeating and bursting ones as in panels B and C; see also Figure S3) for GUVs composed of POPC containing increasing molar fraction of POPG symmetrically and asymmetrically distributed in the membrane leaflets obtained via inverted emulsion method (solid and half-filled squares) and electroformation (solid circles). For GUVs obtained via inverted emulsion approach, average values and standard deviations for measurements on 4 to 6 vesicle preparations per composition are shown (more than 10 vesicles per preparation were monitored). For electroformed GUVs, average values and standard deviations for a number of measurements are shown for 1 vesicle preparation per composition (around 10 vesicles per composition were monitored). Measurements were made in the presence of 0.1 mM EDTA (except for 5 mol% PG sGUVs and aGUVo, prepared via the inverted emulsion protocol. Multiple GUV samples were prepared for each condition and used for the electroporation experiments (n ≥ 3).

We then explored the destabilization fraction *X_dest_* for asymmetric GUVs comparing it to that of symmetric GUVs, whereby both were prepared via the inverted emulsion technique. Interestingly, already small charge asymmetries of 5 mol% POPG considerably enhanced the membrane destabilization and were independent of the direction of the asymmetry (aGUVi or aGUVo), Figure 4D. Therefore, we conclude that charge asymmetry indeed plays an important role in membrane destabilization, rendering the membranes more prone to disturbance events and less able to fully reseal even at low molar fractions of charged species.

To explore the origin of vesicle destabilization, we measured the pore edge tension in these membranes. This parameter reflects the work performed to expand the pore boundary by a unit length and is dependent on membrane composition. The edge tension was obtained from the relaxation dynamics of macropore closure, following an approach reported earlier ^29^ and using an automated image analysis methodology ^34^, see Materials and Methods and Figure S5 for example measurements. Since a significant difference in *X_dest_* was observed between sGUVs prepared by electroformation or the inverted emulsion protocol, we first compared edge tension values of these two systems. Interestingly, there was no difference in pore edge tension γ when comparing sGUVs of the same composition (pure POPC or POPC with 10 mol% POPG) prepared by both methods (Figure S6). Hence, the observed increased destabilization of vesicles prepared by inverted emulsion was not related to hindering macropore closure in the membrane. We conclude that the specific membrane composition, and in particular, the presence of oil residues, affects the intrinsic membrane response towards destabilization, but the oil molecules are not edge active and thus do not influence the measured edge tension values.

Figure 5 shows the edge tension data measured for aGUVs with increasing molar fraction of POPG asymmetry (green circles and red squares). These results are a compilation of data for aGUVo and aGUVi with and without EDTA, known to remove possible calcium ions as contaminants from the medium, which can bind to PG lipids when present at low concentration and alter their properties^32, 33, 75^. No significant differences were observed for the same fraction of POPG irrespective of the leaflet location or the presence of EDTA (see Figure S7). Data previously measured for sGUVs (grown by electroformation) with the same total fraction of POPG are also shown in Figure 5 for comparison (black data, obtained from Lira et al. ^33^). The mean values with standard deviations for all conditions are shown in Table 1. Inverted emulsion GUVs made of pure POPC (0 mol% POPG) have edge tension comparable to literature data ^29, 33, 76, 77^. However, aGUVs containing 5, 10 and 17.5 mol% POPG showed a significant edge tension reduction. Comparing vesicles with the same surface charge composition of one of the leaflets, aGUVs containing 25 mol% POPG showed even stronger reduction in the edge tension (∼15 pN) compared to sGUVs that expose the same POPG fraction but contain twice higher amount of anionic lipid in total (∼23 pN for 50 mol% in total) ^33, 34^, see Table 1.

**Figure 5.**
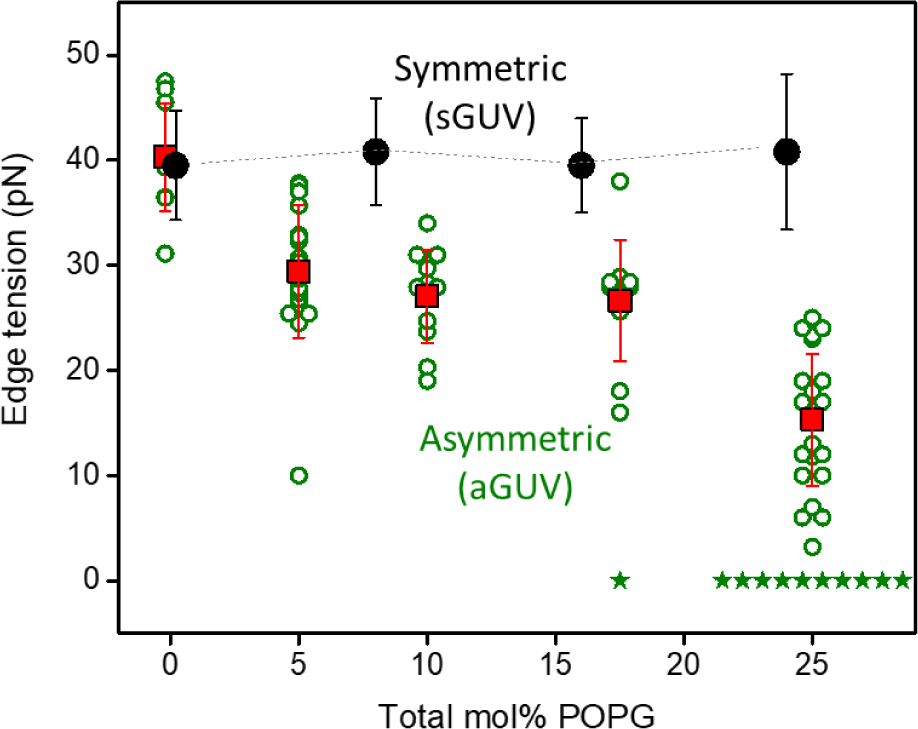
Pore edge tension values of GUVs composed of POPC containing increasing molar fractions of POPG, that were either symmetrically (black solid circles) or asymmetrically distributed (green open circles and red squares). The pore edge tension values for sGUVs are obtained from vesicles produced via electroformation (data published in Lira et al. ^33^). The open green circles represent measurements of individual vesicles. Green stars indicate vesicles that burst after the pulse and are indicated as having a value of the edge tension close to 0. The data for pure POPC membranes (0 mol% POPG) are slightly offset in composition for visibility. Since the type of membrane asymmetry (in terms of POPG location in the outer or inner leaflet) did not affect the pore edge tension, the data from aGUVi and aGUVo, with and without EDTA was combined (see also Figure S7, where data from aGUVi and aGUVo are given separately). A total of 87 vesicles were measured (10 to 34 vesicles per membrane composition). Multiple GUV samples were prepared for each condition and used for the electroporation experiments (n ≥ 3).

**Table 1.**
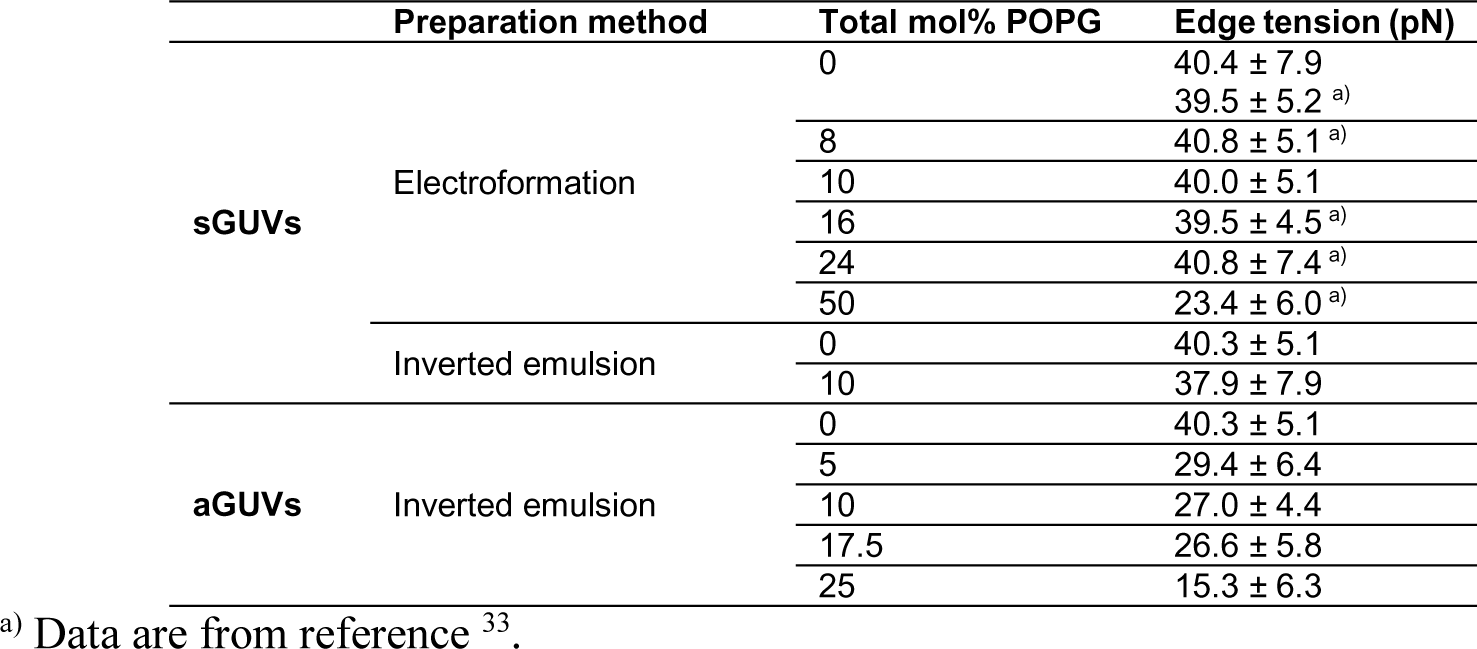
Edge tension values measured for symmetric (sGUVs) and for asymmetric (aGUVs) of POPC with increasing total mole fraction of POPG.

The edge tension results show that the increase in the molar fraction of POPG results in membranes that are more prone to poration (less energy is needed for pore expansion), and this trend was significantly enhanced when the charges were asymmetrically distributed. A much lower fraction of the charged lipid in the asymmetric membrane compared to the symmetric one is sufficient to cause significant reduction in the edge tension values. When discussing results on sGUVs and aGUVs, we made the comparison based on the total amount of POPG that can either be homogeneously distributed between both monolayers (sGUVs) or almost entirely restricted to one of the monolayers (aGUVs). We also considered the comparison based not on the total POPG amount but on the POPG fraction in the POPG-rich leaflet (in this case, the amount of POPG in the POPG-rich leaflet of asymmetric membranes is almost double the one in the symmetric ones with the same total POPG fraction), because it could be that the POPG-rich leaflet dictates the behavior of the whole bilayer. These analyses for the fraction of destabilized GUVs, *X_dest_*, and edge tension are provided in Figure S8. Even when the symmetric membrane has as much POPG as the POPG-rich side of the asymmetric one, aGUVs are still more unstable than sGUVs, both regarding *X_dest_* and pore edge tension.

Above we observed that the preparation method had no effect on the edge tension of symmetric membranes. On the other hand, the fraction of destabilized vesicles, *X_dest_*, which quantifies the occurrence of leaky membranes and vesicle burst after electroporation, was significantly higher for vesicles prepared by the inverted emulsion protocol, even in symmetric cases. Therefore, we hypothesized that traces of oil used to disperse the lipids from the two different monolayers that are present in the membrane are affecting membrane stability. Presumably, the oil confined between the membrane leaflets can influence their degree of interdigitation and coupling, thus affecting the overall stability of the membrane. However, this speculation remains to be explored and the precise amount of oil quantified. To confirm that the main source of instability was brought by charge asymmetry, we also generated membrane asymmetry in the GUVs by a very different approach, namely by varying the pH across the membrane.

### Charge asymmetry caused by different pH values across the membrane also destabilizes membranes

The phosphate group of POPG has a pKa around 4 ^78^. To generate asymmetry, we prepared sGUVs via electroformation containing 20 mol% POPG at neutral pH and then dispersed the GUVs in low pH solution, so that protonation of POPG in the outer layer generated charge asymmetry across the membrane. Figure 6A shows the fraction of destabilized GUVs after electroporation (*X_dest_*) for sGUVs made of POPC with 20 mol% POPG prepared in neutral pH and then dispersed in solutions of different lower pH values down to pH 3; note that the data corresponding to conditions of pH 7 represents the behavior of symmetric membranes (sGUVs). The scatter in the data is somewhat larger compared to that observed for asymmetric membranes prepared with the inverted emulsion method. We speculate that it could be caused by permeation of the hydronium ions across the membrane decreasing the degree of asymmetry. Despite this concern, the data show that *X_dest_* increases significantly for pH 3, which is below the pKa of the phosphate group, but does not change considerably when the external pH is between 4 and 7 (see blue data points in Figure 6A). As a control, we also prepared GUVs made of pure POPC and of POPC with 50 mol% POPG and measured *X_dest_* when the external solution was neutral or at pH 3. As expected, POPC was not destabilized in any pH, because the pKa of its phosphate group is lower than 3 ^79, 80^ (black data in Figure 6A), whereas POPC with 50 mol% POPG showed already a significant destabilization at neutral pH, as reported earlier ^33^, but even stronger when at pH 3 (red data in Figure 6A).

**Figure 6.**
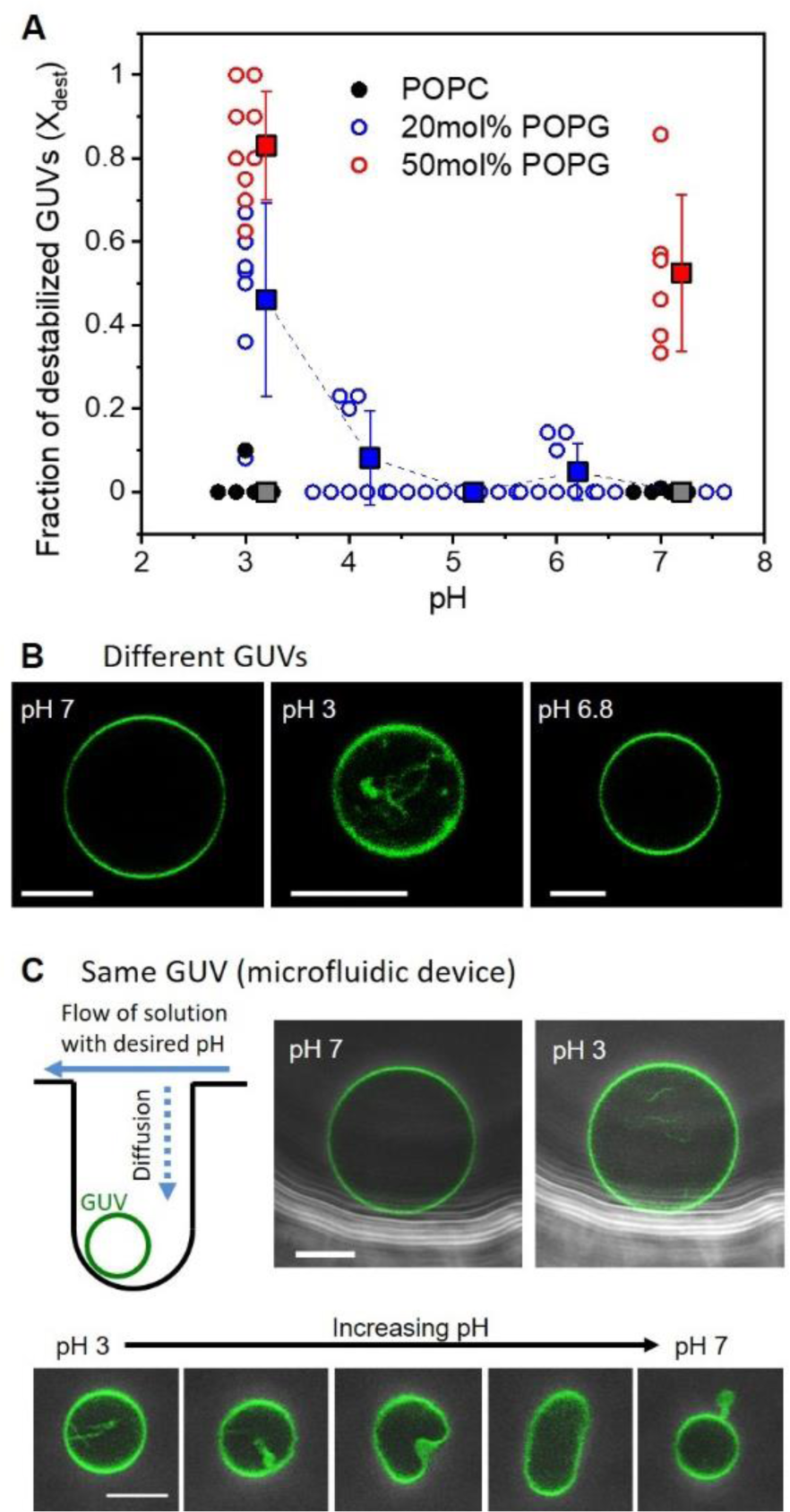
Membrane charge asymmetry generated by different pH across the membrane destabilizes GUVs. (A) Fraction of destabilized vesicles as a function of the external pH of GUVs grown in neutral pH (internal pH 7; note that data acquired at pH 7 correspond to symmetric sGUVs). Circles show values for each observation chamber and filled squares (shifted to the right for clarity) represent the mean values with standard deviation. (B) Representative confocal microscopy images of GUVs of POPC with 20 mol% POPG (0.5 mol% PG-NBD) obtained after dispersing the GUVs of the same sample in pH 7, pH 3 and back to pH 6.8. Scale bars represent 10 μm. Multiple GUV samples were prepared for each condition and used for the electroporation experiments (n ≥ 3). (C) Single-vesicle experiment demonstrating membrane asymmetry generated by exchanging the solution outside the GUV, using a microfluidic trapping device (see also Figure S9). As shown in the sketch, a GUV is trapped in a dead-end channel oriented perpendicular to the flow of the solution that is exchanged. In the dead-end channel, the GUV is shielded from the perturbation of the hydrodynamic flow, which allows to follow the same vesicle over time. Overlay of phase contrast and confocal microscopy images of GUVs of POPC with 20 mol% POPG (1 mol% PG-NBD) trapped in the dead-end channels of the microfluidic device. The outer solution is exchanged with solutions of desired pH. Upon exchanging the solution from pH 7 to pH 3, the vesicle develops inward tube (upper couple of images). Increasing the pH back to 7, causes a vesicle with internal tubules to expel the excess membrane area into outward bud (lower set of images; the time between the first and last image is approximately 5 min).

In summary, the destabilization effect of charged lipids in asymmetric membranes was already pronounced at low POPG fraction as a result of extremely reduced pore edge tension, whereas anionic sGUVs were only affected at membrane compositions of around 50 mol% POPG. We speculate that one important destabilization factor, in addition to the effect of charged lipids, is a high spontaneous curvature that is caused by the area mismatch between the leaflets of membranes with increased POPG asymmetry.

The main concern about this approach is whether the pH asymmetry is indeed maintained during the observation and whether the protonated POPG flips across the membrane, thus obliterating the asymmetry ^81^. Since POPG and its analogue counterpart NBD-PG are symmetrically distributed, the quenching assay is of no use here, and in addition, sodium dithionite is much more unstable at acidic pH. Another way to qualitatively confirm the presence of membrane asymmetry is to probe for spontaneous curvature changes expressed in tubulation of GUVs with excess area ^59–61^. Since protonated POPG has a smaller area per headgroup than its charged state ^82^, charge asymmetry will also cause area imbalance giving rise to membrane spontaneous curvature. We noticed that after dispersion of GUVs of POPC with 20 mol% POPG in solutions of pH 3, inward tubes were detected in some GUVs, whereas when no pH asymmetry was present, GUVs were mainly spherical with smooth membrane and free of tubes (see Figure 6B). To show that the asymmetry was maintained after dispersing the GUVs in pH 3, we increased again the pH of the external solution after inner tubes were formed at pH 3. The tubes were suppressed, showing that the area of the external layer was reestablished and brought back to that of the internal one, and no flip-flop of the protonated PG occurred.

These bulk experiment although encouraging, suffer from the disadvantage that the prehistory of the individual vesicles selected for observation is unknown. To confirm that asymmetry is established, we performed single-vesicle experiments where an individual sGUV with internal pH of 7 is exposed to a solution of pH 3 under constant microscopy monitoring. The vesicles were trapped using a microfluidic trap described previously^67^. As expected, and upon exchange of their external solution, the GUVs responded by forming inward-pointing tubes stabilized by the negative spontaneous curvature (see Figure 6C), confirming the induced pH asymmetry for POPC vesicles with 20 mol% POPG. Conversely, increasing the external pH causes inward tubes to be suppressed (Figure 6C). No tubes were generated in GUVs made of pure POPC.

## Conclusions

Model membranes such as GUVs can be versatile tools to understand the importance and influence of asymmetry in membranes. To draw the right conclusions, the membranes need to be well characterized. A useful approach to verify membrane asymmetry is the quenching assay, although its application requires precaution. We showed that the stabilization of the dithionite ion and radical is crucial for a successful quenching assay. We identified important conditions that have to be considered and tested for each sample system, to avoid any misinterpretation of obtained results. As described by McIntyre and Sleight, sodium dithionite has to be dissolved in a buffer of alkaline pH ^62^, and, most importantly, should be prepared always immediately before usage. The destabilization of the quenching agent and thereby formation of undesired decomposition products can be reduced by usage of low concentrated stock solutions. To avoid membrane destabilization by the presence of dithionite decomposition products (e.g. hydrogen sulphite and hydrogen sulphate), the dithionite concentration should be adjusted to the minimal quenching concentration and subsequent dilution is required after quenching of the outer leaflet is finished. Despite the multiple reported protocols for quenching assays performed on SUVs, LUVs, GUVs and cells ^59, 62–64^, the needed precaution for the handling of sodium dithionite is often underrated, which makes it difficult for readers to reproduce the experimental protocol. Moreover, different membrane systems can show different membrane permeability of the quenching agent ^63, 64^. Our work should raise awareness of the demanding character of the used quenching agent and provide a guide to optimize and adjust the quenching conditions for different samples and experimental setups.

We then investigated membrane stability upon electroporation as a function of charge asymmetry. The results presented here emphasize the impact of anionic lipids on the stability of model membranes, in which the charge distribution is closer to the reality of the cell membrane. We considered not only the effect of increasing the fraction of charged lipids but also the charge lipid asymmetry existing between the membrane leaflets. The latter was established in two ways – using the inverted emulsion protocol for GUV preparation and as induced by pH asymmetry in the solutions across the membrane. As discussed in the Introduction, leaflet asymmetry is capable of dictating various cellular functions by modifying the material properties of membranes and although some studies have investigated lipid asymmetry among monolayers, none of them directly addressed the effect of charge asymmetry between them. To our knowledge, this is the first work to show that charge asymmetry plays an important role in cell membrane destabilization and permeability after electroporation. The origin of this destabilization is partially related to changes in membrane composition as reflected in the changes in the edge tension values. This result can be juxtaposed to data showing changes in the bending rigidity of asymmetric *vs* symmetric membranes for the same overall membrane composition ^66^. This finding was recently interpreted as potentially arising from differential stress (resulting not from compositional but area difference of the leaflets ^73, 83^). It remains to be shown whether membrane destabilization as demonstrated here is a result of such differential stress. We expect that our finding for this asymmetry-enhanced destabilization will contribute towards understanding of the arsenal of recovery and pore-resealing mechanisms developed by cells in wound healing processes.

Finally, our results also suggested that the inverted emulsion method for generating asymmetric GUVs adds some instability to the membrane that is still to be investigated but should be considered when studying membrane parameters that can be affected by a possible oil contamination. The presence of oil in the membrane could affect the degree of interleaflet interdigitation. Molecular dynamic simulations in this direction could shed light in this direction.

## Associated Content

Supporting Information is available free of charge at https://pubs.acs.org/doi…

Data demonstrating the importance of pH of the sodium dithionite stock solution; dependence of the quenching efficiency on the incubation time and quencher concentration; quantification of the destabilizing effects caused by electric pulses; examples of pore closure dynamics and edge tension measurements for symmetrical and asymmetrical GUVs; comparison of edge tension measurements on symmetrical GUVs prepared by electroformation and inverted emulsion technique and by orientation of charge asymmetry and presence of EDTA; fraction of destabilized vesicles as a function of molar fraction of POPG in the POPG-rich side; single-vesicle experiments with pH exchange.

## Author information

### Author contributions

RD and KAR proposed and supervised the project. All authors designed the experiments, FL and MS performed the experiments with bulk GUV studies and analyzed the data. SP performed the microfluidic single-vesicle experiments. FL, MS, RD and KR wrote the manuscript. All authors edited the manuscript. Fernanda Leomil and Mareike Stephan contributed equally to this work.

### Notes

The authors declare no conflict of interest

## Acknowledgements

MS acknowledges funding from the International Max Planck Research School on Multiscale BioSystems. SP was supported by the Max Planck School for Matter to Life.

## References

(1) Lorent, J. H.; Levental, K. R.; Ganesan, L.; Rivera-Longsworth, G.; Sezgin, E.; Doktorova, M.; Lyman, E.; Levental, I. Plasma Membranes Are Asymmetric in Lipid Unsaturation, Packing and Protein Shape. Nat. Chem. Biol. 2020, 16 (6), 644–652. 10.1038/s41589-020-0529-6.

(2) Bretscher, M. S. Asymmetrical Lipid Bilayer Structure for Biological Membranes. Nat. New Biol. 1972 23661 1972, 236 (61), 11–12. 10.1038/newbio236011a0.

(3) Devaux, P. F. Static and Dynamic Lipid Asymmetry in Cell Membranes. Biochemistry 1991, 30 (5), 1163–1173. 10.1021/bi00219a001.

(4) Fadok, V. A.; Henson, P. M. Apoptosis: Getting Rid of the Bodies. Curr. Biol. 1998, 8 (19), R693–R695. 10.1016/S0960-9822(98)70438-5.

(5) Lentz, B. R. Exposure of Platelet Membrane Phosphatidylserine Regulates Blood Coagulation. Progress in Lipid Research. September 2003, pp 423–438. 10.1016/S0163-7827(03)00025-0.

(6) Riedl, S.; Rinner, B.; Asslaber, M.; Schaider, H.; Walzer, S.; Novak, A.; Lohner, K.; Zweytick, D. In Search of a Novel Target - Phosphatidylserine Exposed by Non-Apoptotic Tumor Cells and Metastases of Malignancies with Poor Treatment Efficacy. Biochim. Biophys. Acta - Biomembr. 2011, 1808 (11), 2638–2645. 10.1016/j.bbamem.2011.07.026.

(7) Bevers, E. M.; Williamson, P. L. Getting to the Outer Leaflet: Physiology of Phosphatidylserine Exposure at the Plasma Membrane. Physiol. Rev. 2016, 96 (2), 605–645. 10.1152/physrev.00020.2015.

(8) Fadok, V. A.; Bratton, D. L.; Frasch, S. C.; Warner, M. L.; Henson, P. M. The Role of Phosphatidylserine in Recognition of Apoptotic Cells by Phagocytes. Cell Death and Differentiation. Nature Publishing Group July 1998, pp 551–562. 10.1038/sj.cdd.4400404.

(9) Callahan, M. K.; Williamson, P.; Schlegel, R. A. Surface Expression of Phosphatidylserine on Macrophages Is Required for Phagocytosis of Apoptotic Thymocytes. Cell Death Differ. 2000 77 2000, 7 (7), 645–653. 10.1038/sj.cdd.4400690.

(10) Daleke, D. L. Phospholipid Flippases. J. Biol. Chem. 2007, 282 (2), 821–825. 10.1074/JBC.R600035200.

(11) van Meer, G. Dynamic Transbilayer Lipid Asymmetry. Cold Spring Harb. Perspect. Biol. 2011, 3 (5), 1–11. 10.1101/CSHPERSPECT.A004671.

(12) Allhusen, J. S.; Conboy, J. C. The Ins and Outs of Lipid Flip-Flop. Acc. Chem. Res. 2017, 50 (1), 58–65. 10.1021/acs.accounts.6b00435.

(13) Doktorova, M.; Symons, J. L.; Levental, I. Structural and Functional Consequences of Reversible Lipid Asymmetry in Living Membranes. Nat. Chem. Biol. 2020, 16 (12), 1321–1330. 10.1038/s41589-020-00688-0.

(14) McNeil, P. L.; Steinhardt, R. A. Loss, Restoration, and Maintenance of Plasma Membrane Integrity. J. Cell Biol. 1997, 137 (1), 1. 10.1083/JCB.137.1.1.

(15) McNeil, P. L.; Steinhardt, R. A. Plasma Membrane Disruption: Repair, Prevention, Adaptation. Annu. Rev. Cell Dev. Biol. 2003, 19, 697–731. 10.1146/ANNUREV.CELLBIO.19.111301.140101.

(16) Kinosita, K.; Ashikawa, I.; Saita, N.; Yoshimura, H.; Itoh, H.; Nagayama, K.; Ikegami, A. Electroporation of Cell Membrane Visualized under a Pulsed-Laser Fluorescence Microscope. Biophys. J. 1988, 53 (6), 1015–1019. 10.1016/S0006-3495(88)83181-3.

(17) Kinosita, K.; Tsong, T. Y. Voltage-Induced Pore Formation and Hemolysis of Human Erythrocytes. Biochim. Biophys. Acta - Biomembr. 1977, 471 (2), 227–242. 10.1016/0005-2736(77)90252-8.

(18) Mir, L. M.; Orlowski, S.; Belehradek, J.; Paoletti, C. Electrochemotherapy Potentiation of Antitumour Effect of Bleomycin by Local Electric Pulses. Eur. J. Cancer Clin. Oncol. 1991, 27 (1), 68–72. 10.1016/0277-5379(91)90064-K.

(19) Belehradek, M.; Domenge, C.; Luboinski, B.; Orlowski, S.; Belehradek, J.; Mir, L. M. Electrochemotherapy, a New Antitumor Treatment. First Clinical Phase I-II Trial. Cancer 1993, 72 (12), 3694–3700. 10.1002/1097-0142(19931215)72:12<3694::AID-CNCR2820721222>3.0.CO;2-2.

(20) Mali, B.; Jarm, T.; Snoj, M.; Sersa, G.; Miklavcic, D. Antitumor Effectiveness of Electrochemotherapy: A Systematic Review and Meta-Analysis. Eur. J. Surg. Oncol. 2013, 39 (1), 4–16. 10.1016/j.ejso.2012.08.016.

(21) Frandsen, S. K.; Vissing, M.; Gehl, J. A Comprehensive Review of Calcium Electroporation—A Novel Cancer Treatment Modality. Cancers (Basel). 2020, 12 (2), 290. 10.3390/cancers12020290.

(22) Yarmush, M. L.; Golberg, A.; Serša, G.; Kotnik, T.; Miklavčič, D. Electroporation-Based Technologies for Medicine: Principles, Applications, and Challenges. Annu. Rev. Biomed. Eng. 2014, 16 (1), 295–320. 10.1146/annurev-bioeng-071813-104622.

(23) Lambricht, L.; Lopes, A.; Kos, S.; Sersa, G.; Préat, V.; Vandermeulen, G. Clinical Potential of Electroporation for Gene Therapy and DNA Vaccine Delivery. Expert Opin. Drug Deliv. 2016, 13 (2), 295–310. 10.1517/17425247.2016.1121990.

(24) Keating, A.; Toneguzzo, F. Gene Transfer by Electroporation: A Model for Gene Therapy. Prog. Clin. Biol. Res. 1990, 333, 491–498.

(25) Rols, M. P. Electropermeabilization, a Physical Method for the Delivery of Therapeutic Molecules into Cells. Biochim. Biophys. Acta - Biomembr. 2006, 1758 (3), 423–428. 10.1016/J.BBAMEM.2006.01.005.

(26) Dimova, R.; Marques, C. The Giant Vesicle Book; CRC Press: Boca Raton, 2019. 10.1201/9781315152516.

(27) Riske, K. A.; Dimova, R. Electro-Deformation and Poration of Giant Vesicles Viewed with High Temporal Resolution. Biophys. J. 2005, 88 (2), 1143–1155. 10.1529/biophysj.104.050310.

(28) Riske, K. A.; Dimova, R. Electric Pulses Induce Cylindrical Deformations on Giant Vesicles in Salt Solutions. Biophys. J. 2006, 91 (5), 1778–1786. 10.1529/biophysj.106.081620.

(29) Portet, T.; Dimova, R. A New Method for Measuring Edge Tensions and Stability of Lipid Bilayers: Effect of Membrane Composition. Biophys. J. 2010, 99 (10), 3264–3273. 10.1016/j.bpj.2010.09.032.

(30) Sabri, E.; Aleksanyan, M.; Brosseau, C.; Dimova, R. Effects of Solution Conductivity on Macropore Size Dynamics in Electroporated Lipid Vesicle Membranes. Bioelectrochemistry 2022, 147, 108222. 10.1016/j.bioelechem.2022.108222.

(31) Portet, T.; Mauroy, C.; Démery, V.; Houles, T.; Escoffre, J.-M.; Dean, D. S.; Rols, M.-P. Destabilizing Giant Vesicles with Electric Fields: An Overview of Current Applications. J. Membr. Biol. 2012, 245 (9), 555–564. 10.1007/s00232-012-9467-x.

(32) Riske, K. A.; Knorr, R. L.; Dimova, R. Bursting of Charged Multicomponent Vesicles Subjected to Electric Pulses. Soft Matter 2009, 5 (10), 1983. 10.1039/b900548j.

(33) Lira, R. B.; Leomil, F. S. C.; Melo, R. J.; Riske, K. A.; Dimova, R. To Close or to Collapse: The Role of Charges on Membrane Stability upon Pore Formation. Adv. Sci. 2021, 8 (11), 2004068. 10.1002/advs.202004068.

(34) Leomil, F. S. C.; Zoccoler, M.; Dimova, R.; Riske, K. A. PoET: Automated Approach for Measuring Pore Edge Tension in Giant Unilamellar Vesicles. Bioinforma. Adv. 2021, 1 (1). 10.1093/bioadv/vbab037.

(35) Aleksanyan, M.; Lira, R. B.; Steinkühler, J.; Dimova, R. GM1 Asymmetry in the Membrane Stabilizes Pores. Biophys. J. 2022, 121 (17), 3295–3302. 10.1016/j.bpj.2022.06.011.

(36) Brochard-Wyart, F.; de Gennes, P. G.; Sandre, O. Transient Pores in Stretched Vesicles: Role of Leak-Out. Phys. A Stat. Mech. its Appl. 2000, 278 (1–2), 32–51. 10.1016/S0378-4371(99)00559-2.

(37) Angelova, M. I.; Soléau, S.; Méléard, P.; Faucon, F.; Bothorel, P. Preparation of Giant Vesicles by External AC Electric Fields. Kinetics and Applications. In Trends in Colloid and Interface Science VI; Helm, C., Lösche, M., Möhwald, H., Eds.; Steinkopff: Darmstadt, 1992; pp 127–131.

(38) Cheng, H.-T.; Megha; London, E. Preparation and Properties of Asymmetric Vesicles That Mimic Cell Membranes. J. Biol. Chem. 2009, 284 (10), 6079–6092. 10.1074/jbc.M806077200.

(39) Cheng, H.-T.; London, E. Preparation and Properties of Asymmetric Large Unilamellar Vesicles: Interleaflet Coupling in Asymmetric Vesicles Is Dependent on Temperature but Not Curvature. Biophys. J. 2011, 100 (11), 2671–2678. 10.1016/j.bpj.2011.04.048.

(40) Visco, I.; Chiantia, S.; Schwille, P. Asymmetric Supported Lipid Bilayer Formation via Methyl-β-Cyclodextrin Mediated Lipid Exchange: Influence of Asymmetry on Lipid Dynamics and Phase Behavior. Langmuir 2014, 30 (25), 7475–7484. 10.1021/la500468r.

(41) Pautot, S.; Frisken, B. J.; Weitz, D. A. Engineering Asymmetric Vesicles. Proc. Natl. Acad. Sci. U. S. A. 2003, 100 (19), 10718–10721. 10.1073/pnas.1931005100.

(42) Pautot, S.; Frisken, B. J.; Weitz, D. A. Production of Unilamellar Vesicles Using an Inverted Emulsion. Langmuir 2003, 19 (7), 2870–2879. 10.1021/la026100v.

(43) Moga, A.; Yandrapalli, N.; Dimova, R.; Robinson, T. Optimization of the Inverted Emulsion Method for High-Yield Production of Biomimetic Giant Unilamellar Vesicles. ChemBioChem 2019, 20 (20), 2674–2682. 10.1002/cbic.201900529.

(44) Noireaux, V.; Libchaber, A. A Vesicle Bioreactor as a Step toward an Artificial Cell Assembly. Proc. Natl. Acad. Sci. 2004, 101 (51), 17669–17674. 10.1073/pnas.0408236101.

(45) Nishimura, K.; Suzuki, H.; Toyota, T.; Yomo, T. Size Control of Giant Unilamellar Vesicles Prepared from Inverted Emulsion Droplets. J. Colloid Interface Sci. 2012, 376 (1), 119–125. 10.1016/j.jcis.2012.02.029.

(46) Stachowiak, J. C.; Richmond, D. L.; Li, T. H.; Liu, A. P.; Parekh, S. H.; Fletcher, D. A. Unilamellar Vesicle Formation and Encapsulation by Microfluidic Jetting. Proc. Natl. Acad. Sci. 2008, 105 (12), 4697–4702. 10.1073/pnas.0710875105.

(47) Abkarian, M.; Loiseau, E.; Massiera, G. Continuous Droplet Interface Crossing Encapsulation (CDICE) for High Throughput Monodisperse Vesicle Design. Soft Matter 2011, 7 (10), 4610–4614. 10.1039/c1sm05239j.

(48) Shum, H. C.; Lee, D.; Yoon, I.; Kodger, T.; Weitz, D. A. Double Emulsion Templated Monodisperse Phospholipid Vesicles. Langmuir 2008, 24 (15), 7651–7653. 10.1021/la801833a.

(49) Enoki, T. A.; Feigenson, G. W. Asymmetric Bilayers by Hemifusion: Method and Leaflet Behaviors. Biophys. J. 2019, 117 (6), 1037–1050. 10.1016/j.bpj.2019.07.054.

(50) Feigenson, G. W.; Huang, J.; Enoki, T. A. An Unexpected Driving Force for Lipid Order Appears in Asymmetric Lipid Bilayers. J. Am. Chem. Soc. 2023, 145 (40), 21717–21722. 10.1021/jacs.3c05081.

(51) Yeagle, P. L.; Hutton, W. C.; Martin, R. B. Transmembrane Asymmetry of Vesicle Lipids. J. Biol. Chem. 1976, 251 (7), 2110–2112. 10.1016/S0021-9258(17)33662-1.

(52) Franzin, C. M.; Macdonald, P. M. Detection and Quantification of Asymmetric Lipid Vesicle Fusion Using Deuterium NMR. Biochemistry 1997, 36 (9), 2360–2370. 10.1021/bi9621270.

(53) Traïkia, M.; Warschawski, D. E.; Lambert, O.; Rigaud, J.-L.; Devaux, P. F. Asymmetrical Membranes and Surface Tension. Biophys. J. 2002, 83 (3), 1443–1454. 10.1016/S0006-3495(02)73915-5.

(54) Marquardt, D.; Heberle, F. A.; Miti, T.; Eicher, B.; London, E.; Katsaras, J.; Pabst, G. 1 H NMR Shows Slow Phospholipid Flip-Flop in Gel and Fluid Bilayers. Langmuir 2017, 33 (15), 3731–3741. 10.1021/acs.langmuir.6b04485.

(55) Gerelli, Y.; Porcar, L.; Lombardi, L.; Fragneto, G. Lipid Exchange and Flip-Flop in Solid Supported Bilayers. Langmuir 2013, 29 (41), 12762–12769. 10.1021/la402708u.

(56) Nguyen, M. H. L.; DiPasquale, M.; Rickeard, B. W.; Doktorova, M.; Heberle, F. A.; Scott, H. L.; Barrera, F. N.; Taylor, G.; Collier, C. P.; Stanley, C. B.; Katsaras, J.; Marquardt, D. Peptide-Induced Lipid Flip-Flop in Asymmetric Liposomes Measured by Small Angle Neutron Scattering. Langmuir 2019, 35 (36), 11735–11744. 10.1021/acs.langmuir.9b01625.

(57) Karamdad, K.; Law, R. V.; Seddon, J. M.; Brooks, N. J.; Ces, O. Studying the Effects of Asymmetry on the Bending Rigidity of Lipid Membranes Formed by Microfluidics. Chem. Commun. 2016, 52 (30), 5277–5280. 10.1039/C5CC10307J.

(58) Lipowsky, R. Spontaneous Tubulation of Membranes and Vesicles Reveals Membrane Tension Generated by Spontaneous Curvature. Faraday Discuss. 2013, 161, 305–331. 10.1039/C2FD20105D.

(59) Steinkühler, J.; De Tillieux, P.; Knorr, R. L.; Lipowsky, R.; Dimova, R. Charged Giant Unilamellar Vesicles Prepared by Electroformation Exhibit Nanotubes and Transbilayer Lipid Asymmetry. Sci. Rep. 2018, 8 (1), 1–9. 10.1038/s41598-018-30286-z.

(60) Dasgupta, R.; Miettinen, M. S.; Fricke, N.; Lipowsky, R.; Dimova, R. The Glycolipid GM1 Reshapes Asymmetric Biomembranes and Giant Vesicles by Curvature Generation. Proc. Natl. Acad. Sci. 2018, 115 (22), 5756–5761. 10.1073/pnas.1722320115.

(61) Karimi, M.; Steinkühler, J.; Roy, D.; Dasgupta, R.; Lipowsky, R.; Dimova, R. Asymmetric Ionic Conditions Generate Large Membrane Curvatures. Nano Lett. 2018, 18 (12), 7816–7821. 10.1021/acs.nanolett.8b03584.

(62) McIntyre, J. C.; Sleight, R. G. Fluorescence Assay for Phospholipid Membrane Asymmetry. Biochemistry 1991, 30 (51), 11819–11827. 10.1021/bi00115a012.

(63) Pomorski, T.; Herrmann, A.; Zachowski, A.; Devaux, P. F.; Müllery, P. Rapid Determination of the Transbilayer Distribution of NBD-Phospholipids in Erythrocyte Membranes with Dithionite. Mol. Membr. Biol. 1994, 11 (1), 39–44. 10.3109/09687689409161028.

(64) Angeletti, C.; Nichols, J. W. Dithionite Quenching Rate Measurement of the Inside−Outside Membrane Bilayer Distribution of 7-Nitrobenz-2-Oxa-1,3-Diazol-4-Yl-Labeled Phospholipids †. Biochemistry 1998, 37 (43), 15114–15119. 10.1021/bi9810104.

(65) Angelova, M. I.; Dimitrov, D. S. Liposome Electroformation. Faraday Discuss. Chem. Soc. 1986, 81 (0), 303. 10.1039/dc9868100303.

(66) Elani, Y.; Purushothaman, S.; Booth, P. J.; Seddon, J. M.; Brooks, N. J.; Law, R. V.; Ces, O. Measurements of the Effect of Membrane Asymmetry on the Mechanical Properties of Lipid Bilayers. Chem. Commun. 2015, 51 (32), 6976–6979. 10.1039/C5CC00712G.

(67) Pramanik, S.; Steinkühler, J.; Dimova, R.; Spatz, J.; Lipowsky, R. Binding of His-Tagged Fluorophores to Lipid Bilayers of Giant Vesicles. Soft Matter 2022, 18 (34), 6372–6383. 10.1039/D2SM00915C.

(68) Lem, W. J.; Wayman, M. Decomposition of Aqueous Dithionite. Part I. Kinetics of Decomposition of Aqueous Sodium Dithionite. Can. J. Chem. 1970, 48 (5), 776–781. 10.1139/v70-126.

(69) Vegunta, V. L. A Study on the Thermal Stability of Sodium Dithionite Using ATR-FTIR Spectroscopy. 2016, No. June, 272.

(70) Burlamacchi, L.; Guarini, G.; Tiezzi, E. Mechanism of Decomposition of Sodium Dithionite in Aqueous Solution. Trans. Faraday Soc. 1969, 65, 496–502. 10.1039/TF9696500496.

(71) Kamiya, K.; Kawano, R.; Osaki, T.; Akiyoshi, K.; Takeuchi, S. Cell-Sized Asymmetric Lipid Vesicles Facilitate the Investigation of Asymmetric Membranes. Nat. Chem. 2016, 8 (9), 881–889. 10.1038/nchem.2537.

(72) Stephan, M. S.; Dunsing, V.; Pramanik, S.; Chiantia, S.; Barbirz, S.; Robinson, T.; Dimova, R. Biomimetic Asymmetric Bacterial Membranes Incorporating Lipopolysaccharides. Biophys. J. 2023. 10.1016/j.bpj.2022.12.017.

(73) Hossein, A.; Deserno, M. Spontaneous Curvature, Differential Stress, and Bending Modulus of Asymmetric Lipid Membranes. Biophys. J. 2020, 118 (3), 624–642. 10.1016/j.bpj.2019.11.3398.

(74) Bhatia, T.; Christ, S.; Steinkühler, J.; Dimova, R.; Lipowsky, R. Simple Sugars Shape Giant Vesicles into Multispheres with Many Membrane Necks. Soft Matter 2020, 16 (5), 1246–1258. 10.1039/C9SM01890E.

(75) Riske, K. A.; Döbereiner, H.-G.; Lamy-Freund, M. T. Comment on “Gel-Fluid Transition in Dilute versus Concentrated DMPG Aqueous Dispersions.” J. Phys. Chem. B 2003, 107 (22), 5391–5392. 10.1021/jp027077p.

(76) Mattei, B.; Lira, R. B.; Perez, K. R.; Riske, K. A. Membrane Permeabilization Induced by Triton X-100: The Role of Membrane Phase State and Edge Tension. Chem. Phys. Lipids 2017, 202, 28–37. 10.1016/j.chemphyslip.2016.11.009.

(77) Casadei, B. R.; Dimova, R.; Riske, K. A. Bending Modulus and Edge Tension of Giant Unilamellar Vesicles (GUVS) Composed of Lipid Extracts from Erythrocytes Membranes. Biophys. J. 2018, 114 (3), 94a. 10.1016/j.bpj.2017.11.558.

(78) Watts, A.; Harlos, K.; Maschke, W.; Marsh, D. Control of the Structure and Fluidity of Phosphatidylglycerol Bilayers by PH Titration. Biochim. Biophys. Acta - Biomembr. 1978, 510 (1), 63–74. 10.1016/0005-2736(78)90130-X.

(79) Garidel, P.; Johann, C.; Mennicke, L.; Blume, A. The Mixing Behavior of Pseudobinary Phosphatidylcholine-Phosphatidylglycerol Mixtures as a Function of PH and Chain Length. Eur. Biophys. J. 1997, 26 (6), 447–459. 10.1007/s002490050099.

(80) Moncelli, M. R.; Becucci, L.; Guidelli, R. The Intrinsic PKa Values for Phosphatidylcholine, Phosphatidylethanolamine, and Phosphatidylserine in Monolayers Deposited on Mercury Electrodes. Biophys. J. 1994, 66 (6), 1969–1980. 10.1016/S0006-3495(94)80990-7.

(81) Khalifat, N.; Rahimi, M.; Bitbol, A.-F.; Seigneuret, M.; Fournier, J.-B.; Puff, N.; Arroyo, M.; Angelova, M. I. Interplay of Packing and Flip-Flop in Local Bilayer Deformation. How Phosphatidylglycerol Could Rescue Mitochondrial Function in a Cardiolipin-Deficient Yeast Mutant. Biophys. J. 2014, 107 (4), 879–890. 10.1016/j.bpj.2014.07.015.

(82) Khalifat, N.; Puff, N.; Bonneau, S.; Fournier, J.-B.; Angelova, M. I. Membrane Deformation under Local PH Gradient: Mimicking Mitochondrial Cristae Dynamics. Biophys. J. 2008, 95 (10), 4924–4933. 10.1529/biophysj.108.136077.

(83) Lyman, E.; Sodt, A. J. Differential or Curvature Stress? Modus Vivendi. Biophys. J. 2020, 118 (3), 535–537. 10.1016/j.bpj.2019.11.3399.

