## Supporting information for "Bilayer charge asymmetry and oil residues destabilize membranes upon poration"

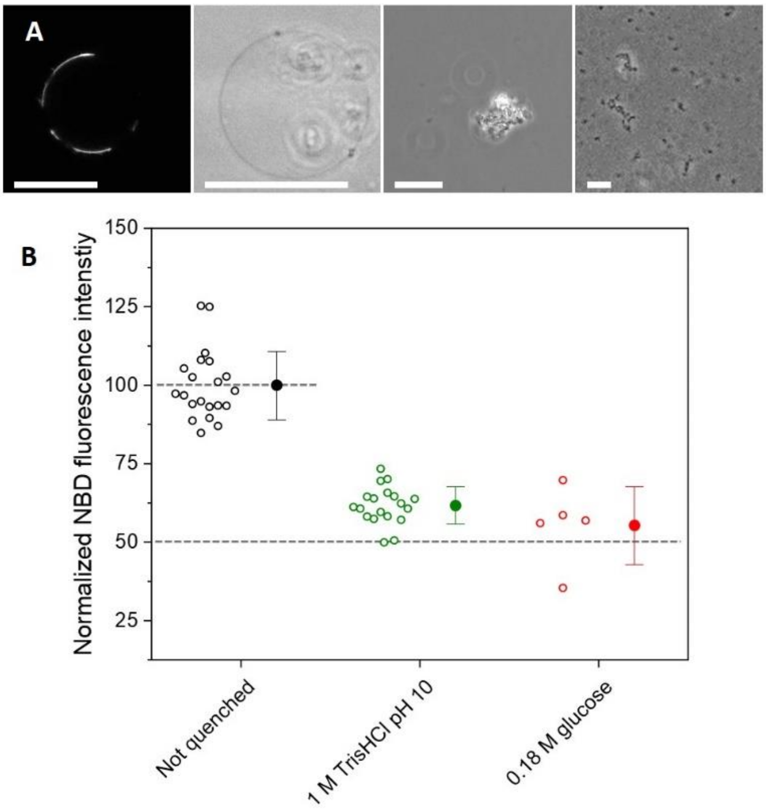


**Figure S1**: Non-buffered dithionite solutions affect GUV stability and do not allow for consistent quenching experiments. All experiments were conducted with symmetric and electroformed GUVs. (A) Examples of GUVs and lipid aggregates observed when diluting POPC GUVs (1 mol % NBD-PG) with 0.1 M $Na_2S_2O_4$ prepared in 0.18 M glucose to a final concentration of 2.5 mM $Na_2S_2O_4$. The first image was acquired with confocal and the others with phase contrast microscopy. The scalebars correspond to 25 µm. (B) Normalized fluorescence intensities before quenching (black circles) and in the presence of 2.5 mM $Na_2S_2O_4$ added from stock solutions of 0.1 M dithionite prepared in 1 M TrisHCl pH 10 (green circles) or in 0.18 M glucose (red circles). Each open symbol represents measurements on one vesicle and mean values with standard deviation are shown on the right. The dashed lines are guides to the eye and indicate the original vesicles intensity (unquenched) and half of this initial mean value. The scatter in the data is a result of the different vesicles sizes (i.e. equatorial sections located higher above the chamber bottom and deeper in the sample) and inhomogeneous distribution of the dye across vesicles. Therefore, the data should be interpreted with precaution and quantitate comparison should be considered only from samples quenched at the same conditions. Note that the data in Figure 2B for the case of the non-buffered conditions (0.18 M glucose) is scarce because of the small number and defective GUVs surviving the treatment.

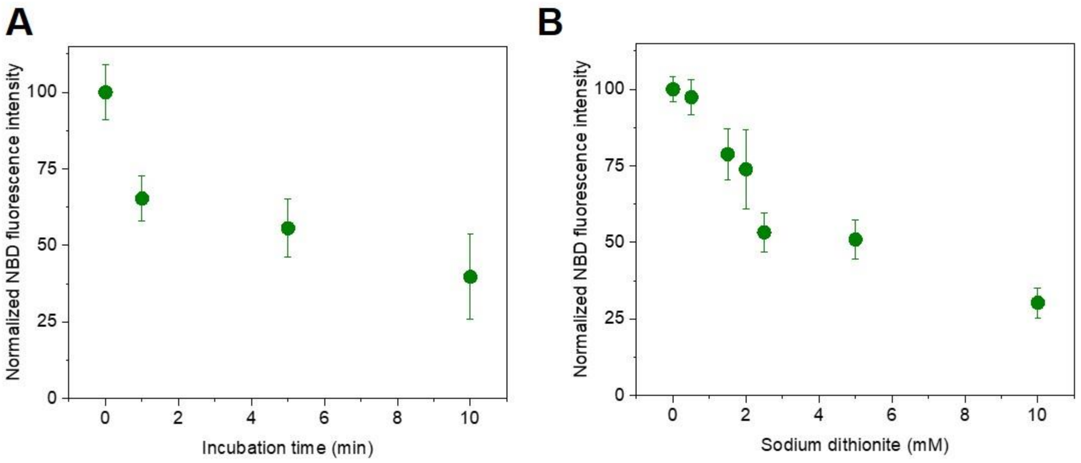


**Figure S2.** (A) Normalized fluorescence intensities before (not quenched, indicated as 0 incubation time) and after quenching with 2.5 mM $Na_2S_2O_4$ for different incubation times followed by 5-fold dilution – same data as in Figure 3C but with linear x-coordinate. (B) Normalized fluorescence intensities before (0) and after quenching of different sodium dithionite concentration for 5 minutes incubation time followed by 5-fold dilution – same data as in Figure 3D but with linear x-coordinate. All experiments were conducted with symmetric and electroformed GUVs.

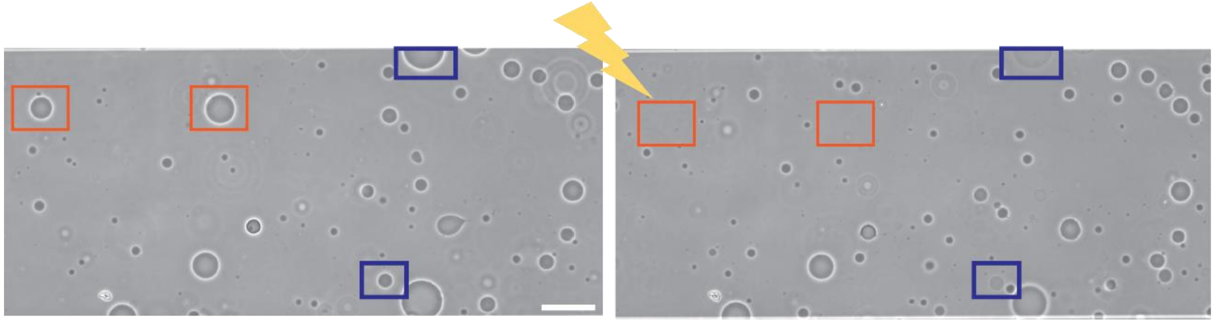

**Figure S3.** Quantifying the number of GUVs that undergo bursting or contrast loss. Phase contrast snapshots of a GUV sample in the electroporation chamber before (left) and after (right) pulse application and assessing the fraction of destabilized vesicles ($X_{dest}$). Upon poration, GUVs marked by orange rectangles bursted ($n_{burst}$) and those marked by blue rectangles became leaky ($n_{perm}$). $X_{dest}$ is calculated as $X_{dest} = (n_{burst} + n_{perm})/n_{GUVs}$, where $n_{GUVs}$ is the total number of GUVs before pulse application. Scale bar: 100 µm. Sample: symmetric electroformed POPC:POPG (1:1) GUVs.

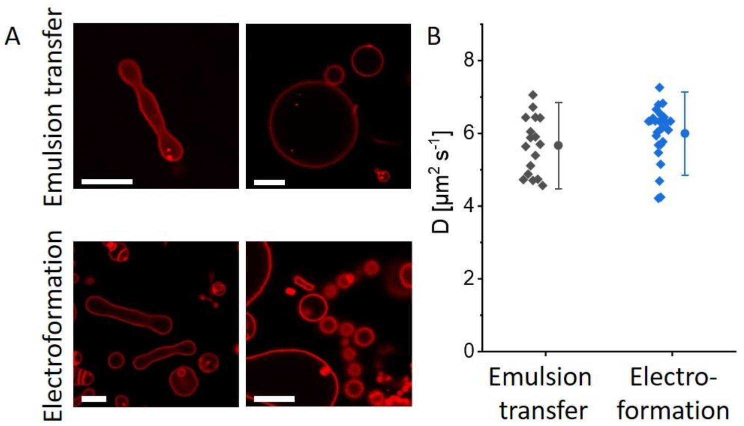


**Figure S4.** Membrane morphology and diffusivity in POPC GUVs prepared using the inverted emulsion protocol and electroformation. **A:** GUVs were deflated by addition of hypertonic glucose solution (1 M glucose). The inner sucrose solution had an osmolarity of 650 mOsmol and the osmolarity of the outer solution was increased to final 800 mOsmol. In bothsamples the deflation resulted in prolate or multisphere GUVs as expected for vesicles with sucrose/glucose asymmetry across the membrane[1]. The scale bars correspond to 10 µm. **B:** Lipid diffusion coefficient (measured with TexasRed-DHPE) obtained from fluorescence recovery after photobleaching. The measured diffusion coefficients are not significantly different from each other. Data adapted from reference[2] M. S. Stephan, V. Dunsing, S. Pramanik, S. Chiantia, S. Barbirz, T. Robinson, R. Dimova, Biomimetic asymmetric bacterial membranes incorporating lipopolysaccharides, Biophys. J. 2023, Copyright (2023), with permission from Elsevier.

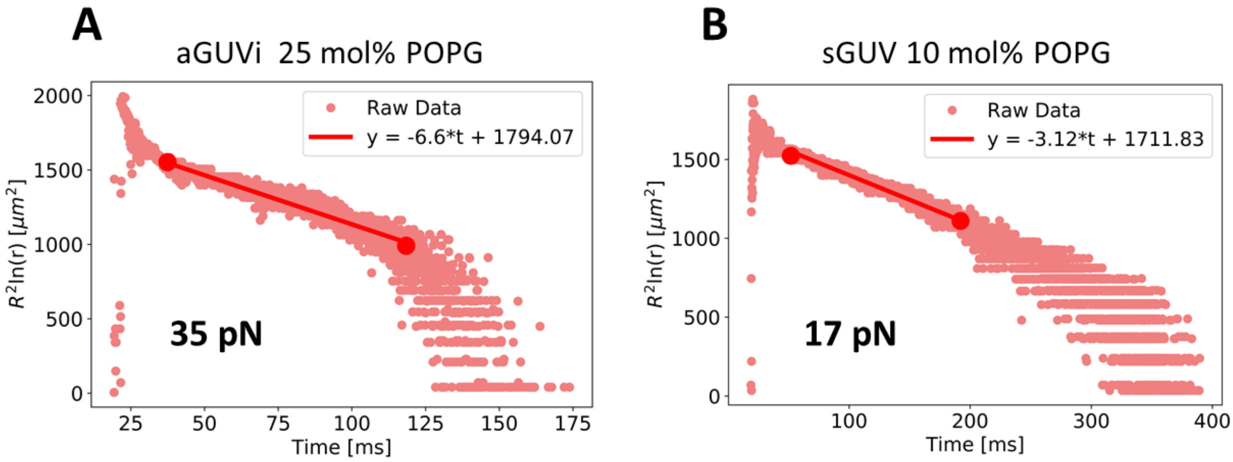


**Figure S5.** Example measurements of the pore edge tension. The vesicles are subjected to a porating pulse (3 kV/cm, 150 µs), see Materials and Methods section for details. The radius of a formed macropore, *r*, in a vesicle of radius, *R*, is detected using the home-developed software PoET [3]. The pore dynamics follows four well-defined stages; (i) quick opening, (ii) maximum size stage, (iii) slow closure, limited by leak-out, and (iv) rapid closure. According to a theoretical model developed in [4,5], the slow pore closure (third stage) can be directly related to the edge tension through $R^2 ln(r) = -(2\gamma/3\pi\eta)t + C$, where $\eta$ is the medium viscosity, $t$ is time and $C$ is a constant. The edge tension $\gamma$ is directly calculated from the slope of the linear dependence of $R^2 ln(r)$ with time. The two examples presented here correspond to pore edge tension measurements for (A) sGUV 10 mol% POPG and (B) aGUVi 25 mol% POPG. In the graphs, the pore size $r$ in µm is rescaled by $l$ = 1 µm to avoid taking the logarithm of dimensional value. The respective edge tension values obtained from the slopes of the fits (red solid lines) are 35 pN and 17 pN.

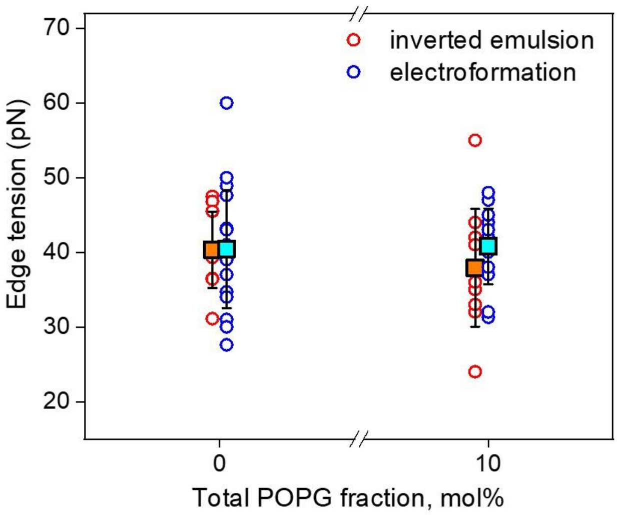


**Figure S6.** Comparison of pore edge tension values for POPC and POPC with 10 mol% POPG GUVs prepared via inverted emulsion (red circles) and electroformation (blue circles). Each point represents a measurement on one single vesicle and squares show the mean values with standard deviation. A total of 54 vesicles were measured, 12 to 18 vesicles per membrane composition.

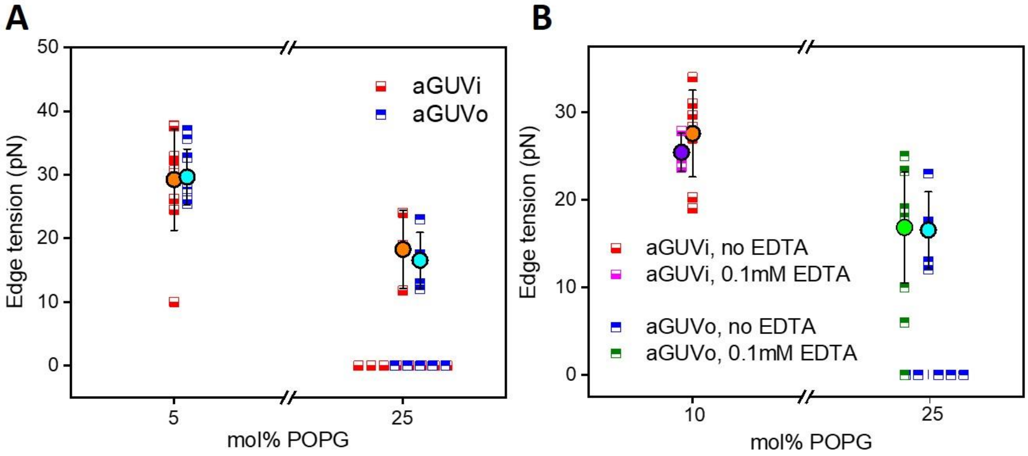


**Figure S7.** Location of charge in asymmetric membranes in GUVs prepared via the inverted emulsion technique and presence of EDTA does not influence edge tension values. (A) Asymmetric GUVs containing 5 and 25 mol% POPG in the outer leaflet (aGUVo) and in the inner leaflet (aGUVi). (B) Asymmetric GUVs containing 10 and 25 mol% PG in aGUVi and aGUVo in the absence and in the presence of 0.1 mM EDTA. Each half-filled square represents a measurement on one vesicle, mean and standard deviation are shown in circles. Data showing zero edge tension values correspond to GUVs which have burst.

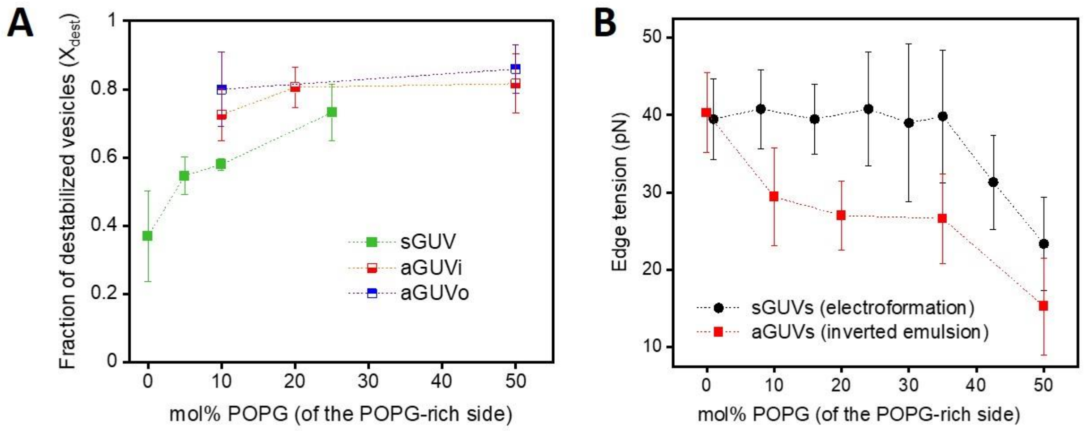


**Figure S8.** Comparison of the (A) fraction of destabilized vesicles and (B) pore edge tension mean values for sGUVs and aGUVs as a function of the molar fraction of POPG in the POPG-rich leaflet. In (A) the sGUVs were also obtained by the inverted emulsion protocol whereas in (B) the sGUVs were grown by electroformation and the data were taken from Lira et al. [6].

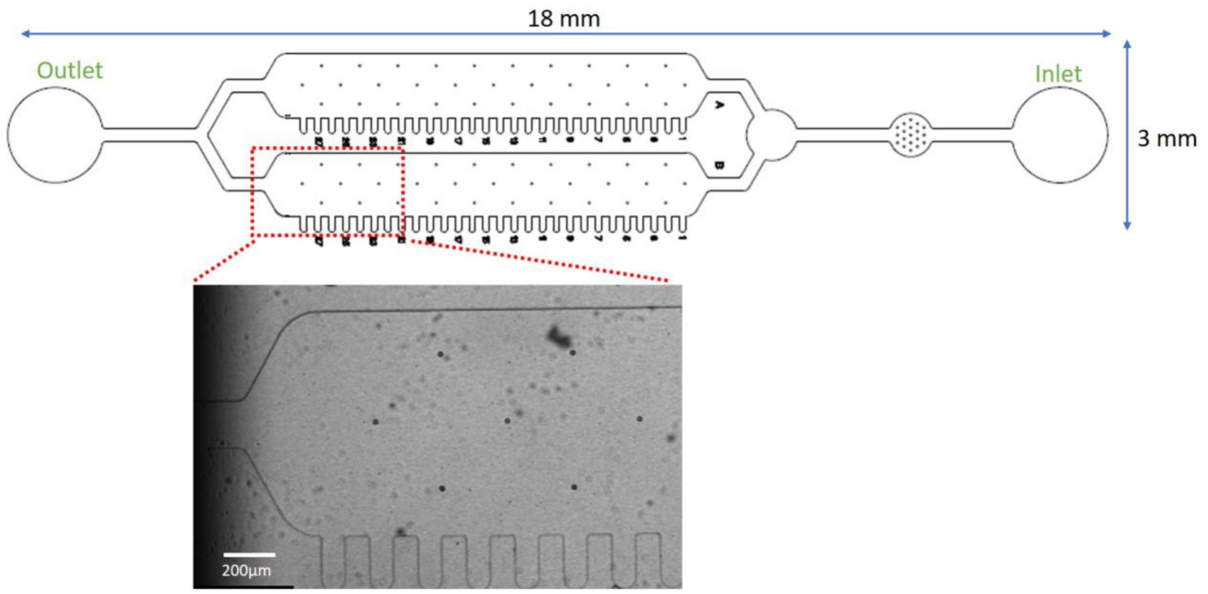


**Figure S9.** Design of the microfluidic chip (full view shown in the lower right corner) with dead-end channels for trapping GUVs and solution exchange[7]. The bright field image at the bottom show a portion of the chip with the side channels.

### Movie captions

**Movie S1.** Vesicle composed of POPC:POPG in the molar ratio of 8:2 and containing 1 mol% NBD-PG was electroformed on ITO glasses in the presence of 200 mM sucrose solution at pH 7. The vesicle was trapped in the microfluidic device and the outer solution was exchanged with an isotonic mixture of glucose and sucrose (in the ratio of 9:1) at pH 7 before starting the recording. During the course of the movie, the outer solution was exchanged with isotonic glucose/sucrose (in the ratio of 9:1) solution at pH 3, resulting in the tubes in the inside of the GUV. The diameter of the GUV was 32 µm. The time stamp corresponds to min:sec.

**Movie S2.** Vesicle composed of POPC:POPG in the molar ratio of 8:2 and containing 1 mol% NBD-PG was electroformed on ITO glasses in the presence of 200 mM sucrose solution at pH 7. The vesicle was trapped in the microfluidic device and the outer solution was exchanged with an isotonic mixture of glucose and sucrose (in the ratio of 9:1) at pH 3 before starting the recording. During the course of the movie, the outer solution was exchanged with an isotonic glucose/sucrose (in the ratio of 9:1) solution at pH 7, resulting in the tubes being pulled out of the GUV and resulting in outward buds and protrusions. The diameter of the GUV was 20 µm. The time stamp corresponds to min:sec.

**References**


[1] T. Bhatia, S. Christ, J. Steinkühler, R. Dimova, R. Lipowsky, *Soft Matter* **2020**, *16*, 1246.

[2] M. S. Stephan, V. Dunsing, S. Pramanik, S. Chiantia, S. Barbirz, T. Robinson, R. Dimova, *Biophys. J.* **2023**, DOI 10.1016/j.bpj.2022.12.017.

[3] F. S. C. Leomil, M. Zoccoler, R. Dimova, K. A. Riske, *Bioinforma. Adv.* **2021**, *1*, DOI 10.1093/bioadv/vbab037.

[4] F. Brochard-Wyart, P. G. de Gennes, O. Sandre, *Phys. A Stat. Mech. its Appl.* **2000**, *278*, 32.

[5] R. Ryham, I. Berezovik, F. S. Cohen, *Biophys. J.* **2011**, *101*, 2929.

[6] R. B. Lira, F. S. C. Leomil, R. J. Melo, K. A. Riske, R. Dimova, *Adv. Sci.* **2021**, *8*, 2004068.

[7] S. Pramanik, J. Steinkühler, R. Dimova, J. Spatz, R. Lipowsky, *Soft Matter* **2022**, *18*, 6372.
